# Trehalose-6-phosphate synthase promotes thermotolerance by governing glycolytic flux in *Cryptococcus deneoformans*

**DOI:** 10.64898/2026.07.15.738695

**Authors:** Vikas Yadav, Kahlia A. Carl, Joseph Heitman, Erica J. Washington

## Abstract

Growth at physiologically relevant temperatures is essential for fungal pathogenesis and is controlled by several cellular factors. The evolution of fungal thermotolerance is concerning as warming environments may promote the emergence of new pathogens. Trehalose, a disaccharide absent in mammals, plays a central role in thermotolerance by stabilizing proteins and membranes during heat stress. Trehalose is synthesized from glucose-6-phosphate (G6P) and uridine-diphosphate-glucose (UDPG) in two steps catalyzed by trehalose-6-phosphate synthase (Tps1) and trehalose-6-phosphate phosphatase (Tps2). Here, we investigated genetic suppression of Tps1 function in *Cryptococcus deneoformans*, a species in the *Cryptococcus* pathogenic species complex. Tps1 is essential for growth at 37°C in *C. deneoformans* and spontaneous suppressor mutations restored the growth of *tps1*Δ mutants at 37°C. Whole-genome sequencing followed by variant calling analysis primarily identified loss-of-function mutations in the gene encoding hexokinase 1 (Hxk1). Targeted gene deletion mutants further showed that loss of either *HXK1* or *HXK2* can bypass *tps1*Δ in a carbon source-dependent manner. The *tps1*Δ mutant exhibited elevated hexokinase activity, accumulation of G6P and glycogen, and ATP depletion after heat shock. Deletion of *HXK1* or *HXK2* restored hexokinase activity and partially restored G6P and ATP levels in the *tps1*Δ mutant, while glycogen remained elevated, indicating that excess glycolytic flux underlies the *tps1*Δ high-temperature growth defect. Overall, our study uncovers a previously unappreciated mechanism of Tps1-mediated heat adaptation in *C. deneoformans*, by revealing that Tps1 functions as a critical metabolic gatekeeper that safeguards glycolytic flux to sustain growth at elevated temperatures.

**Article summary:** Trehalose is crucial for fungal thermal adaptation and mutants lacking trehalose are inviable at 37°C. This study examined how genetic suppressors restore viability at 37°C in mutants lacking *TPS1*, which encodes trehalose-6-phosphate synthase. Through whole-genome sequencing of spontaneous suppressor isolates and variant calling analysis, mutations were identified in *HXK1*. Gene deletion mutants and biochemical assays show these mutations alter glycolytic flux. We show that *tps1*Δ mutants exhibit unbridled glycolysis, and their growth at 37°C is restored by *hxk1*Δ mutations that reduce glycolytic flux. This study highlights the interdependence between Tps1 and Hxk1, which may have broader relevance across organisms.

## Introduction

Fungi exist in diverse ecological niches, including soil, plants and animals. Most fungal species found in these niches are unable to cause diseases in humans given that the human body is a hostile environment consisting of high temperature, limiting nutrients, reactive oxygen species, hypoxic conditions, immune system defenses, and the host microbiota (1, 2). However, some fungi can withstand conditions in the host, as well as thwart human immune defenses, enabling the microbe to colonize the human host. As a result, fungi can cause both superficial and invasive infections, the latter of which result in over 3.75 million deaths worldwide each year (3, 4).

Due to the elevated temperature of human hosts compared to natural environments where fungi reside, one of the stressful conditions that fungi must tolerate while colonizing human hosts is heat stress. The ability to survive at elevated temperatures, called thermotolerance, is a critical factor that allows fungi to cause disease. Fungal adaptation to higher temperatures is alarming given ongoing environmental warming (5–7). The possibility that environmental fungi might evolve to tolerate heat stress and therefore generate new potential pathogens is of grave concern and may have already occurred with the emergence of new fungal pathogens, such as *Candida auris* (6, 8, 9).

*Cryptococcus* spp. are environmental fungi that can cause lethal infections in immunocompromised patients, as well as immunocompetent patients. The human pathogenic *Cryptococcus* species complex accounts for more than 180,000 deaths worldwide each year and approximately 19% of HIV/AIDS-related deaths (10). *Cryptococcus neoformans* (serotype A) is the predominant cause of HIV/AIDS-associated cryptococcal meningoencephalitis (10, 11). Although the greatest burden of disease caused by *C. neoformans* occurs in sub-Saharan Africa, *C. neoformans* has a global distribution (10). In contrast, *Cryptococcus deneoformans* (serotype D) infects nearly 20% of HIV-positive patients in Europe (12, 13). *Cryptococcus deuterogattii*, a representative of the *Cryptococcus gattii* species complex (serotypes B and C), has emerged in the North American Pacific Northwest where it is causing life-threatening disease in immunocompetent hosts (14, 15).

Thermotolerance is required for *Cryptococcus* to cause disease in humans (16, 17). There are several molecular mechanisms that contribute to thermotolerance in *Cryptococcus* including the trehalose biosynthesis pathway (18–21). Trehalose is a nonreducing disaccharide that is composed of two glucose molecules (α-D-glucopyranosyl-[1,1]-α-D-glucopyranoside) that are linked by an α,α-1,1-glycosidic bond. Trehalose serves two critical functions in fungal physiology. The first is to protect proteins and cell membranes from a series of stresses such as dehydration, exposure to salt/high ionic strength, and increases in temperature that occur during human infection (22). Secondly, trehalose is a source of energy. Trehalases cleave the trehalose glycosidic bond to form two molecules of glucose that subsequently enter glycolysis and the pentose phosphate metabolic pathway (22).

The trehalose biosynthesis pathway in *Cryptococcus* consists of two enzymes: trehalose-6-phosphate synthase, Tps1, and trehalose-6-phosphate phosphatase, Tps2. Tps1 utilizes the substrates uridine-diphosphate-glucose (UDP-glucose) and glucose-6-phosphate (G6P) to generate the intermediate, trehalose-6-phosphate (T6P). This substrate-assisted catalysis requires proper alignment of each substrate to form T6P (23). Tps1 is a highly specific enzyme, given that UDP-galactose, the saccharide of which is an epimer of glucose, cannot be converted to T6P by Tps1 (24). Fungal Tps1 enzymes, as well as other described Tps1 enzymes, belong to the GT-B family of retaining glycosyltransferases (25). Members of the GT-B retaining glycosyltransferase family are characterized by two facing Rossman-like domains. The outcome of the reactions catalyzed by these enzymes is an anomeric bond in which the configuration of the product is retained, compared to the donor substrate.

Structures of the *Candida albicans* Tps1 enzyme revealed the protein is divided into two catalytic subdomains, in which the N-terminal subdomain binds the G6P acceptor molecule and the C-terminal subdomain binds the UDP-glucose donor (23). This same fold was seen in the cryo-electron microscopy (cryo-EM) structures of *C*. *neoformans* Tps1, which also revealed that *C. neoformans* Tps1 forms a homotetramer (24). These structures allowed description of the molecular transitions from a substrate-free state to a ligand-bound conformation, which is necessary for catalysis. Intriguingly, the product T6P can be cytotoxic to fungal cells at high concentrations, but is dephosphorylated by Tps2 to form the final product, trehalose (22–25).

Despite significant previous studies on the structure and function of Tps1, the mechanism(s) by which Tps1 contributes to thermotolerance in *Cryptococcus* is not well understood. Here we set out to identify temperature-resistant mutants in the *C. deneoformans tps1*Δ background at elevated temperatures to gain insight into the genetic components involved in Tps1-mediated heat tolerance in *C. deneoformans*. In this genetic screen, we isolated 21 independent suppressors that bypass the requirement of Tps1 for growth at 37°C. Whole-genome sequencing followed by variant calling resulted in the identification of mutations in several genes, including the genes encoding hexokinase enzymes in *C. deneoformans*, particularly *HXK1*. 17 of the 21 suppressors had loss-of-function mutations in *HXK1*.

Further analysis revealed that in the absence of Tps1, and its product T6P, energy-generating processes are uninhibited, leading to hyperaccumulation of G6P and glycogen in addition to depletion of ATP. Thermotolerance was re-established in *C. deneoformans tps1*Δ cells when spontaneous loss-of-function mutations in *HXK1* occurred in *tps1*Δ cells. Deletion of *HXK1* or *HXK2* partially restored G6P and ATP levels in the *tps1*Δ mutant, but glycogen levels remained high, consistent with partial rebalancing of glycolytic flux. These findings indicate that glycolysis is elevated in *tps1*Δ cells and that deletion of *HXK1* or *HXK2* brings glycolytic flux closer to wild-type levels. In summary, our study reveals interesting insights into the connection between Tps1, thermotolerance, and glycolysis, all of which are required for the mammalian pathogenicity of *Cryptococcus deneoformans* strains.

## Materials and methods

### Strains and media

*Cryptococcus deneoformans* (previously known as *C. neoformans* var. *neoformans*) reference strains JEC20**a** and JEC21α were used for experiments (26, 27). The strains were cultured in YPD (1% yeast extract, 2% peptone, and 2% glucose) liquid or YPD agar media at 25°C or 37°C for most experiments. In some cases, YP alone or YP supplemented with 2% galactose (YPGal) was used. Gene deletion mutants for *hxk1*Δ and *hxk2*Δ mutants were generated via homologous recombination with CRISPR following the TRACE protocol (28) and transformants were selected on YPD agar media supplemented with either 100 µg/mL nourseothricin or 200 µg/mL G-418. The strains analyzed in this study are presented in Table S1 and primers employed in this study are presented in Table S2.

### Isolation of *TPS1* suppressors

Strains lacking *TPS1* (JEC20 *tps1*Δ and JEC21 *tps1*Δ) were first streaked to obtain single colonies on YPD media at 25°C. Next, 12 independent single colonies were inoculated for each strain in 5 mL YPD media and grown overnight at 25°C. An equal number of cells, equivalent to 1 OD, were plated on YPD agar plates and incubated at 37°C. The plates were incubated for 4 to 5 days to allow the growth of spontaneous mutant strains that produced colonies at 37°C. From each plate, 4 colonies were further streaked on YPD plate and verified for robust growth. Out of these, only one colony was then used for further analysis as an independent genetic mutant for *tps1*Δ suppression, and a total of 21 suppressors were isolated through this approach.

### Whole-genome sequencing and variant calling

The verified suppressors along with the JEC20 *tps1*Δ and JEC21 *tps1*Δ mutant parental strains were subjected to Illumina whole-genome sequencing and variant calling was performed as described previously (29). Briefly, each strain was grown overnight in 5 mL YPD liquid media to collect cells, which were frozen, and then lyophilized before DNA extraction with the MasterPure yeast DNA purification kit (Biosearch Technologies). The isolated DNA was checked for quality via agarose gel electrophoresis and NanoDrop and was quantified by Qubit, and the samples were submitted to Duke Sequencing and Genomic Technologies core for sequencing. KAPA HyperPrep kits were utilized for library prep and sequencing was performed on the Illumina Novaseq platform.

The sequencing reads obtained were checked for quality with FastQC and mapped to *C. deneoformans* reference JEC21 genome available on FungiDB (https://fungidb.org/fungidb/app) using Bowtie2 with default parameters and allowing every read to map only once. The BAM files containing mapped reads were analyzed for variant identification with Geneious Prime. Single Nucleotide Polymorphism (SNP) events covered with at least 90 reads and present in 90% of covering reads were identified and considered for further analysis. Among these, SNPs that were shared between the suppressor isolates and *tps1*Δ mutants were removed from the analysis. This analysis resulted in the identification of only one SNP in the protein-coding regions per isolate for the majority of the isolates. For isolates that did not present any SNP in their protein-coding regions, SNPs present in the introns were considered. For three isolates, no SNPs in the coding region or introns could be identified and these were analyzed by coverage analysis, which revealed absence of read coverage indicating the loss of *CNH01400* locus in them. Combined together, we were able to assign a genetic change in each isolate with the majority of the isolates harboring only one genetic change. The mutations identified are summarized in Table 1.

**Table 1.**
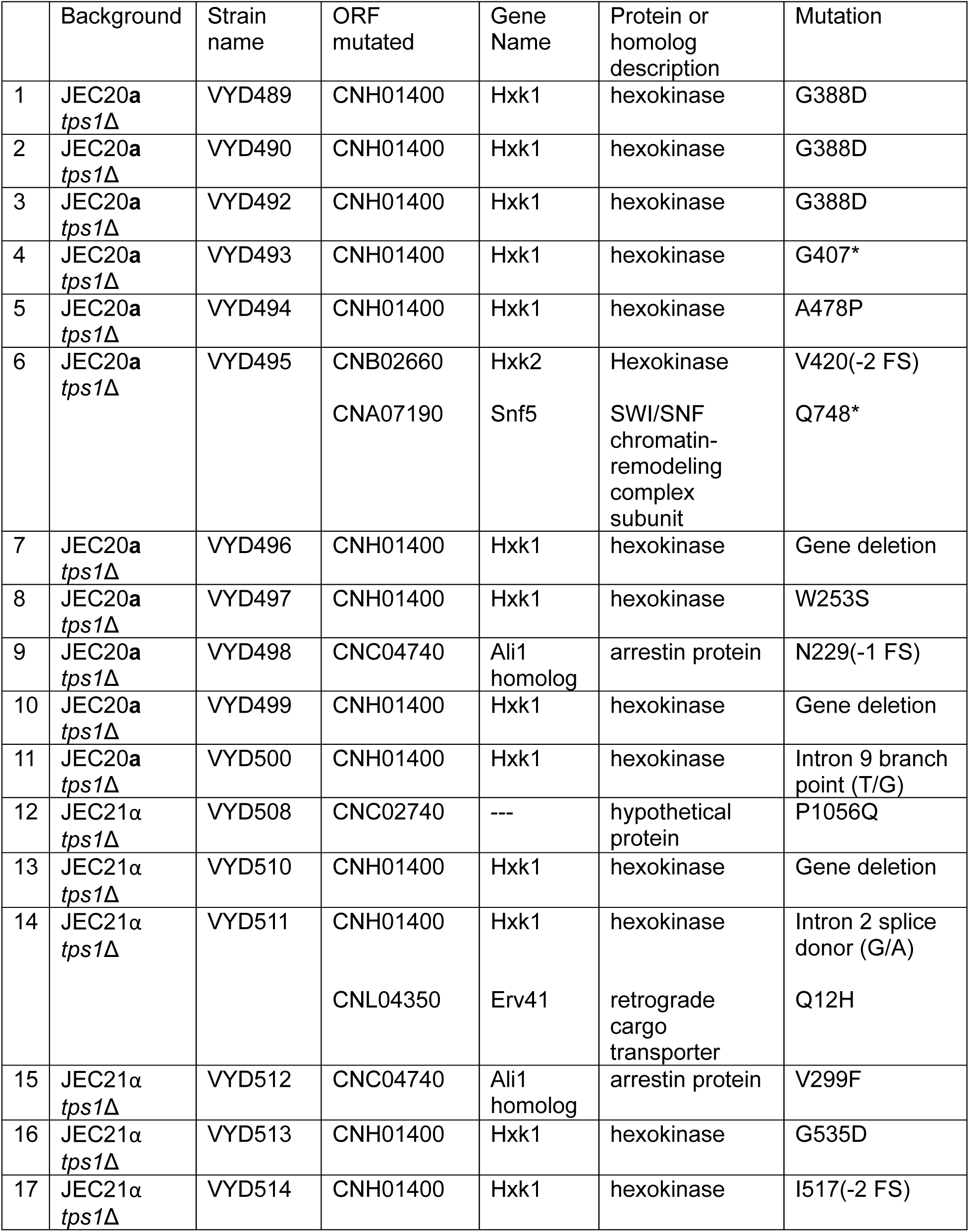
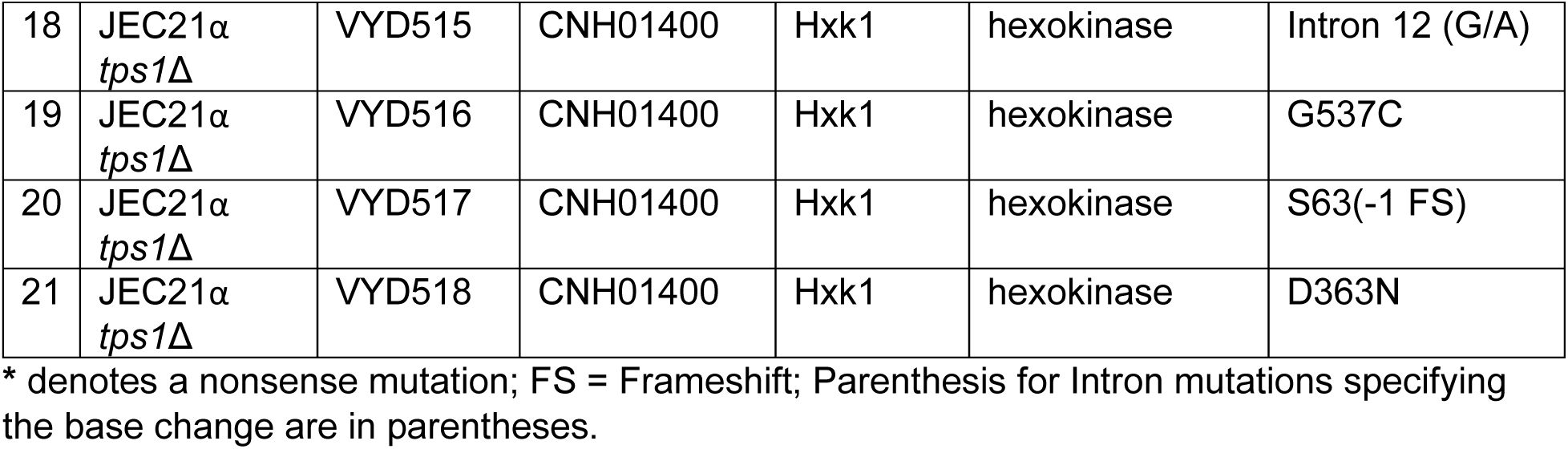

### Serial dilution spotting assays

Strains were grown overnight in 5 mL YPD at 25°C and growth was measured by OD_600_. For each culture, 5 OD equivalent cells were transferred to a fresh tube, centrifuged, washed once with autoclaved dH_2_O and resuspended in 1 mL of dH_2_O to normalize the number of cells across samples. The resuspension was then subjected to 10-fold serial dilution in 96-well plates to generate the dilutions 0, 10^-1^, 10^-2^, 10^-3^, 10^-4^, 10^-5^. From each dilution, 3 µL was spotted onto the YPD, YPGal or YP plates and incubated at 25°C or 37°C for 2-3 days.

### Hexokinase protein purification

The *HXK1* gene from *C. deneoformans* JEC20, lacking the sequence corresponding to the first 96 amino acids, was codon-optimized for expression in *E. coli* (Genscript) and inserted into the pET28a vector, which contains an N-terminal 6xHis-tag with a thrombin cleavage site. Variants of Hxk1 with the point mutations W253S, D290A, G388D, and G537C were generated with the wild-type Hxk1 pET28a plasmid as the template. For protein expression, the constructs were transformed into *Escherichia coli* BL21(DE3) (Life Technologies, Inc.). Cell cultures were grown to an OD_600_ of 0.6 – 0.8 and expression was initiated by the addition of 1 mM isopropyl β-D-1-thiogalactopyranoside (IPTG) at 15°C. The cultures were then allowed to grow for an additional 16 hours prior to harvesting by centrifugation at 4000 rpm at 4°C. After lysis in buffer containing 50 mM Tris-HCl, pH=8.0, 300 mM NaCl, 5 mM MgCl_2_, 5% glycerol, and 5 mM imidazole, supernatants from lysed cultures were loaded onto a HisTrapFF column (Cytiva). The protein was eluted with increasing concentrations of imidazole in the lysis buffer. 5 mL fractions containing 6xHis-Hxk1_97-557_, as determined by SDS-PAGE, were pooled and concentrated using 30K MWCO Amicon Ultra concentrators (Millipore) to 1 mg/mL for downstream applications.

### Hexokinase activity assay

Cells were grown from fresh colonies in 5 mL YPD media to saturation at 25°C with shaking. Cultures were diluted 10-fold and then cells were grown for an additional 8 hours in YPD media at 37°C. The concentration of the cultures was adjusted to 10^7^ cells/mL in sterile YPD. The Cryptococcal cells were washed once with PBS, pelleted by centrifugation for 5 minutes at 4000 rpm, and stored in -80°C in round-bottom culture tubes overnight. The pellet was lyophilized. The pellet was disrupted with the addition of acid-washed glass beads. Samples were disrupted in two 1-minute pulses and kept on ice before and after disruption. The total hexokinase activity was assessed with a colorimetric hexokinase activity assay (Abcam). Absorbance readings were measured at OD_450_ using a kinetic program to measure for 30 minutes. The activity of purified wild-type Hxk1 recombinant protein, as well as the Hxk1 variants with point mutations W253S, D290A, G388D, and G537C, was determined with a colorimetric activity assay (Abcam). Proteins were quantified by Bradford assay (30).

### G6P accumulation assay

Fresh colonies were grown in 5 mL of YPD broth to saturation at either 25°C with shaking. The cultures were diluted 1:10 and then cells were grown for an additional 8 hours in YPD media at either 25°C or 37°C. The concentration of the cultures was adjusted to 10^7^ cells/mL in sterile YPD. Cells were washed once with PBS, pelleted by centrifugation for 5 minutes at 4000 rpm, and stored in -80°C in round-bottom culture tubes overnight. The pellet was lyophilized. The pellet was disrupted with the addition of acid-washed glass beads using two 1-minute pulses and kept on ice before and after disruption. The total G6P was assessed using the colorimetric glucose-6-phosphate assay kit (BioAssay Systems). Absorbance readings were measured at OD_460_ after 20 minutes after the addition of substrate.

### ATP accumulation assay

Cells were grown from fresh colonies in 5 mL YPD media to saturation at 25°C with shaking. Due to the inherently unstable nature of ATP, the cultures were exposed to 37°C for only 1 minute for heat shock and immediately flash frozen in liquid nitrogen. The pellet was disrupted with the addition of acid-washed glass beads. 200 µL of Assay Buffer 23 (Abcam) was added to the disrupted samples prior to cell lysis. Samples were disrupted using two 1-minute pulses and kept on ice before and after disruption. Samples were centrifuged at 10,000 rpm at 4°C for 2 minutes and the supernatants were collected. Cellular ATP was measured with a colorimetric assay with the absorbance measured at OD_570_ (Abcam). Proteins were quantified by Bradford assay (30).

### Glycogen assay with iodine staining

Cells were grown overnight in 5 mL YPD media from fresh colonies. The concentration of the cultures was adjusted to 10^7^ cells/mL in sterile YPD. This stock was then serially diluted 10-fold 5 times. Cells at each of these concentrations were spotted onto YPD agar plates. All plates were incubated at 25°C or 37°C for 48 hours to allow for cell growth. Plates lacking lids were then transferred to petri dishes containing 0.5 grams of evenly distributed iodine crystals. Agar plates were removed from the iodine exposure and then photographed after 8 minutes.

### Glycogen measurement assay

Cells were grown from fresh colonies in 5 mL YPD media to saturation at 25°C with shaking. Cultures were diluted 10-fold and then cells were grown for an additional 8 hours in YPD media at either 25°C or 37°C. 3 mL of cells were collected, wash with cold PBS, and resuspended in 200 µL of sterile water. 100 µL of acid-washed glass beads were added to the samples, which were then disrupted using two 1-minute pulses. Samples were kept on ice before and after disruption, with a 1-minute incubation on ice between pulses. Lysates were collected and centrifuged at 14,000 rpm for 10 minutes at 4°C. The resulting supernatants were boiled for 10 minutes to inactivate enzymatic activity. Protein concentration was then quantified by Bradford assay (30). Glycogen content was measured using the colorimetric Glycogen Assay Kit (Abcam).

Absorbance was measured at OD_570_. To control for background glucose, the mean absorbance of parallel reactions lacking glucoamylase was subtracted from the corresponding reactions containing glucoamylase.

### Statistical analysis

Comparisons between wildtype and variants or mutants were performed using the one-way ANOVA analysis followed by post-hoc Dunnett’s multiple comparisons test. Statistically significant differences are marked with asterisks (* = p <0.05; ** = p < 0.01; *** = p < 0.001; **** = p < 0.0001). Only the p-values for the statistically significant differences compared to the control group are shown.

## Results

### A suppressor screen of a temperature-sensitive *C. deneoformans tps1*Δ mutant

Tps1 is required for thermotolerance (growth at 37°C) of *C. neoformans* (19). We determined that Tps1 is also required for thermotolerance in *C. deneoformans* (Figure 1A). To identify genes involved in Tps1-mediated thermotolerance in *C. deneoformans*, we performed a suppressor screen to isolate spontaneous thermotolerant mutants in a *C. deneoformans tps1*Δ mutant background. JEC20 *tps1*Δ and JEC21 *tps1*Δ strains were plated onto solid YPD medium and incubated at 37°C for 4 to 5 days, at which point spontaneous thermotolerant mutants were observed (Figure 1B). These colonies were further validated for growth at 37°C and only one colony from each replicate culture was selected for analysis. We obtained 11 JEC20 *tps1*Δ and 10 JEC21 *tps1*Δ suppressor mutant colonies with the ability to grow at 37°C (Table 1 and Figure 1C).

**Figure 1.**
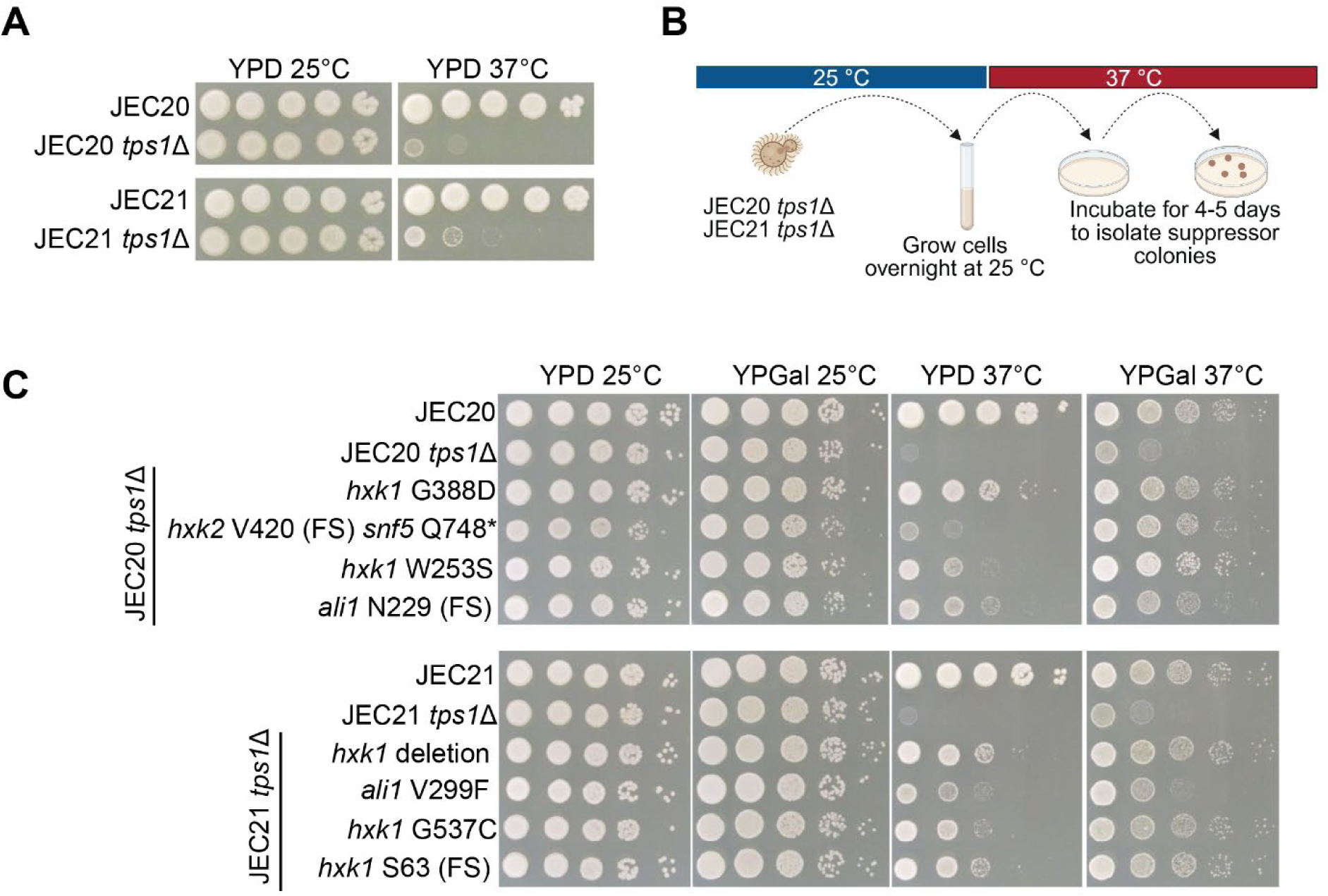
A suppressor screen of the *C. deneoformans tps1*Δ thermosensitive phenotype. **A)** *C. deneoformans* JEC20, *C. deneoformans* JEC20 *tps1*Δ, *C. deneoformans* JEC21, and *C. deneoformans* JEC21 *tps1*Δ serial dilutions were grown on rich yeast-peptone-dextrose (YPD) media at 25°C and 37°C. **B)** Schematic depicting the suppressor screen to isolate suppressor mutations in the *C. deneoformans* JEC20 *tps1*Δ and *C. deneoformans* JEC21 *tps1*Δ thermosensitive background. **C)** Serial dilution assays of suppressors isolated from the screen in either the JEC20 *tps1*Δ or JEC21 *tps1*Δ background grown on YPD and yeast-peptone-galactose (YPGal) at 25°C and 37°C. **\*** denotes a nonsense mutation; FS = Frameshift.

To identify the genetic changes responsible for the suppression of the *tps1*Δ thermosensitive phenotype, we conducted whole-genome sequencing followed by variant calling. Of the 21 *C. deneoformans tps1*Δ thermotolerant suppressors, 17 had mutations in the hexokinase 1 gene, *HXK1* (Table 1). One of the *HXK1* mutants also had a secondary mutation in a gene called *ERV41*. Another suppressor had a mutation in hexokinase 2 (*HXK2*), in combination with a mutation in *SNF5* (Table 1). Other suppressors had mutations in the genes encoding the arrestin protein Ali1 homolog or a hypothetical protein (CNC02740) (Table 1) (31). The suppressor mutants exhibited varying degrees of thermotolerance (Figure 1C).

### Multiple suppressed lines of *C. deneoformans tps1*Δ mutants contain mutations in hexokinase genes

Hexokinase 1 (Hxk1) is a conserved enzyme in plants, animals, and yeasts that phosphorylates intracellular glucose to produce glucose 6-phosphate (G6P) (32). Given that G6P is an important intermediate in glycolysis, glycogen synthesis, and the pentose phosphate pathway, hexokinase 1 initiates the key first step in many energy-generating pathways. However, many hexokinases have dual-functions and serve an additional role in sugar sensing, in the absence of their catalytic function (33). Because our genetic suppressor screen identified multiple independent mutations in *HXK1* gene, we therefore focused on understanding the role of hexokinases in Tps1-dependent thermotolerance.

The *HXK1* mutations identified included stop codon gain mutations, frameshifts, missense mutations, and intron mutations affecting potential splice acceptor/donor/branch sites. Additionally, three isolates harbored substantial gene deletions, leading us to conclude that the suppressor mutations in *HXK1* are loss-of-function mutations (Figure 2A and Table 1). Analysis of deletion flanking regions did not identify any repeat sequences or short homology regions and the exact mechanism causing these deletions is not apparent. The AlphaFold predicted structure of the *C. deneoformans* Hxk1 contains both linked large and small domains, similar to its closest structural homolog, *Kluyveromyces lactis* Hxk1, as well as *C. neoformans* hexokinases (Figure 2B,C and Supplemental Figure 1) (34, 35). There are large and small domains of hexokinases that are required to bind glucose and adenosine triphosphate (ATP) (36–40). The AlphaFold predicted structure of *C. deneoformans* Hxk2 is structurally similar to *C. deneoformans* Hxk1 (Supplemental Figure 1) (35, 41). However, the major difference between the two enzymes is found in the N-terminus. Hxk1 has a 96-residue N-terminal domain that is absent in Hxk2 (Supplemental Figure 1,2).

**Figure 2.**
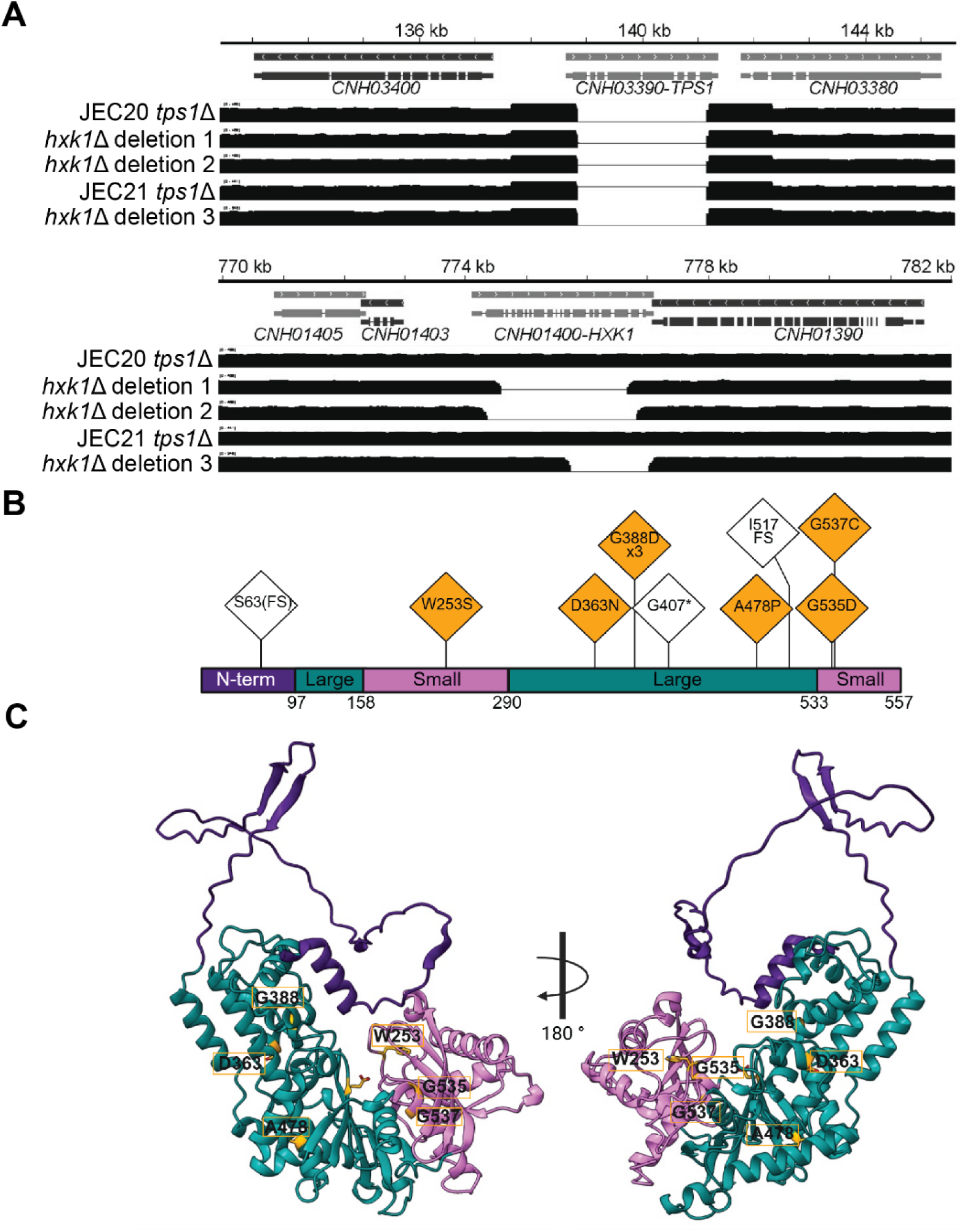
*C. deneoformans tps1*Δ suppressor mutations in Hxk1. **A)** Sequencing analysis revealed precise deletions of the *HXK1* gene occurred in three independent 37°C-temperature-resistant suppressor isolates of *C. deneoformans tps1*Δ mutants. The top panel presents read coverage showing the targeted deletion of *TPS1* gene and the bottom panel depicts the spontaneous deletion events of *HXK1* gene. **B)** Domain organization of the *C. deneoformans* hexokinase Hxk1 with the key domains depicted with the N-terminus in purple, the large domain in teal, and the small domain in pink. Point mutations in Hxk1 identified in the suppressor screen are shown in the orange triangles, depicting the location of the residues in the affected domains. The domain organization is based on the *C. deneoformans* Hxk1 structure predicted by AlphaFold (Q5KCL1) (35, 41). **\*** denotes a nonsense mutation; FS = Frameshift. Produced with Biorender. **C)** Predicted structure, rotated 180°, of *C. deneoformans* Hxk1 (Q5KCL1) as determined by AlphaFold (35, 41). Key domains are colored as described in the previous panel. Key residues identified in suppressor screen are labelled and shown as orange atom-colored stick representations. Generated in ChimeraX.

The Hxk1 point mutations identified in the suppressor strains are located in multiple sites within the predicted structure of the *C. deneoformans* Hxk1 (Figure 2A) and these residues are well-conserved amongst fungal hexokinases (Supplemental Figure 2). For instance, hexokinases are highly conserved amongst most fungal species, with the *Cryptococcus* hexokinases clustering together and indicating that duplication of the hexokinases occurred after speciation (Supplemental Figure 3). Among the identified amino acid mutations, residues W253, G537, and G535 are located in the hexokinase small domain. For instance, W253 is located near the glucose-binding site, while G537 and G535 are found proximal to the ATP-binding sites. D363, A478, and G388 are observed in the hexokinase large domain. G537 and G535 are located in an α-helix that interacts with the large domain. D363, G388, and A478 are not positioned proximal to either substrate-binding site and, therefore, may actually regulate the activity of Hxk1 via an allosteric mechanism. AlphaFold modeling of the *C. deneoformans* Hxk1 variants does not reveal global structural changes compared to the wild-type model (Supplemental Figure 4) (35, 42). *De novo* crystallization, combined with other biochemical approaches, will likely be required to determine how the point mutations affect the structure of the hexokinases.

### Hexokinase point mutations in the *C. deneoformans tps1*Δ background reduce enzyme activity

In this study, we focused on three point mutations in Hxk1 (G537C, W253S, and G388D) occurring in different regions of the Hxk1 protein. Given that hexokinases can function in both glucose metabolism and regulatory, sugar-sensing pathways, we are interested in what the selected point mutations might reveal about these hexokinase functions.

To determine the effect of these point mutations on hexokinase activity, we purified recombinant wild-type 6xHis-Hxk1_97-557_ protein from *E. coli* by affinity chromatography via the hexahistidine tag, along with the G537C, W253S, and G388D variants. We also generated a 6xHis-Hxk1_97-557_ D290A active site mutant as a negative control. This aspartic acid lies in the catalytic center of hexokinase, and the side chain of D290 makes a hydrogen bond with a hydroxyl group of glucose (36). Mutation of this aspartic acid to an alanine or a glycine residue results in a catalytically inactive enzyme (39). The N-terminal region of Hxk1 was not included in the purification construct due to the concern that this large unstructured region might hinder the purification of the recombinant protein. The activity of the Hxk1 enzymes was measured *in vitro* with a colorimetric hexokinase activity assay. Hxk1 G537C and Hxk1 W253D had significant reductions in specific activity compared to wild-type Hxk1, corresponding to approximately 1800-and 14-fold decreases in their specific activity, respectively. Although variants may appear similar in the graphical representation, fold-change values are provided to enable quantitative comparison and to capture subtle but reproducible differences within the dynamic range of the assay. These data indicate that potential disruption of the ATP-binding site and glucose-binding sites in the Hxk1 G537C and Hxk1 W253D variants resulted in significant loss-of-function of Hxk1 (Figure 3A). Hxk1 G388D, the most distal to known substrate-binding pockets, retained the most Hxk1 activity *in vitro*, reducing the activity only 3.4-fold compared to the wild-type Hxk1 enzyme. The Hxk1 D290A variant, which was included as a negative control, decreased the specific activity by approximately 2,300-fold. Taken together, these results are consistent with our hypothesis that Hxk1 mutations found in the suppressor screen result in either complete or partial loss-of-function of Hxk1.

**Figure 3.**
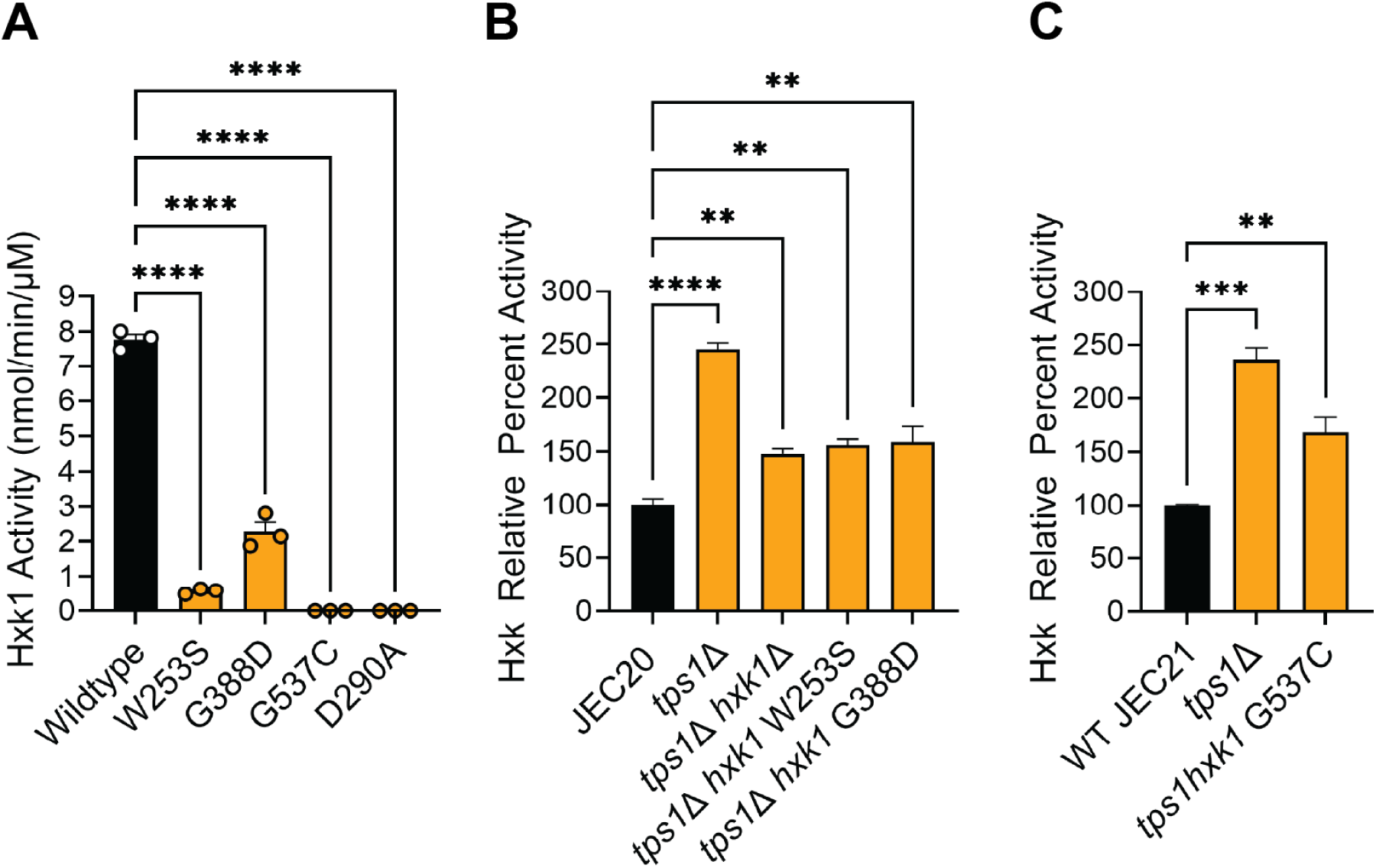
Hexokinase activity in variants of Hxk1 from purified protein and cellular extracts. **A)** Specific activity (nmol/min/µM) of recombinant 6xHis-Hxk1_97-557_ wild-type, 6xHis-Hxk1_97-557_ W253S, 6xHis_97-557_-Hxk1 G388D, 6xHis_97-557_-Hxk1 G537C, and 6xHis_97-557_-Hxk1 D290A. Proteins were expressed in *E. coli* and purified using affinity chromatography. Bars signify the mean ± SEM of technical triplicates. Data shown are representative of three independent experiments. **B,C)** Hexokinase activity in extracts from *C. deneoformans* suppressor strain with the indicated *hxk1* mutation grown on YPD at 37°C. Bars signify the mean ± SEM of biological triplicates. Data shown are representative of three independent experiments.

To determine if these mutations further result in a loss of hexokinase activity in the yeast cells, we measured the cellular hexokinase activity in wild-type yeast cells, as well as an *hxk1*Δ deletion strain and the strains containing Hxk1 point mutations W253S, G388D and G537C in a *tps1*Δ background (Figure 3B,C). Deletion of *TPS1* in both *C. deneoformans* JEC20 and JEC21 backgrounds resulted in a significant 2.4-fold increase in relative hexokinase activity compared to wildtype. Deletion of *HXK1* in the *tps1*Δ background partially reduced hexokinase activity but at 1.5-fold change compared to wild-type levels. The remaining hexokinase activity in the *tps1*Δ *hxk1*Δ strain may be due to the activity of the other hexokinase in *C. deneoformans*, Hxk2. Consistent with the *in vitro* results (Figure 3A), Hxk1 point mutations W253S, G388D, and G537C resulted in significant loss of activity of Hxk1 in extracts from *C. deneoformans* cells (Figure 3B,C).

### Suppressor mutants require specific carbon sources for growth

Hxk1 and Hxk2 intersect with the trehalose biosynthetic pathway (Figure 4A). The first enzyme in the trehalose biosynthesis pathway, Tps1, converts G6P and UDPG to T6P. In *S. cerevisiae* and other fungal organisms, T6P inhibits the activity of Hxk1 and Hxk2, which are required for generating G6P, by phosphorylating glucose (43–46). This tightly controlled pathway links glycolysis and energy-producing pathways with trehalose biosynthesis and thermotolerance. This pathway also indicates the importance of carbon sources for thermotolerance, as G6P is a critical initial component of both glycolysis and trehalose biosynthesis.

**Figure 4.**
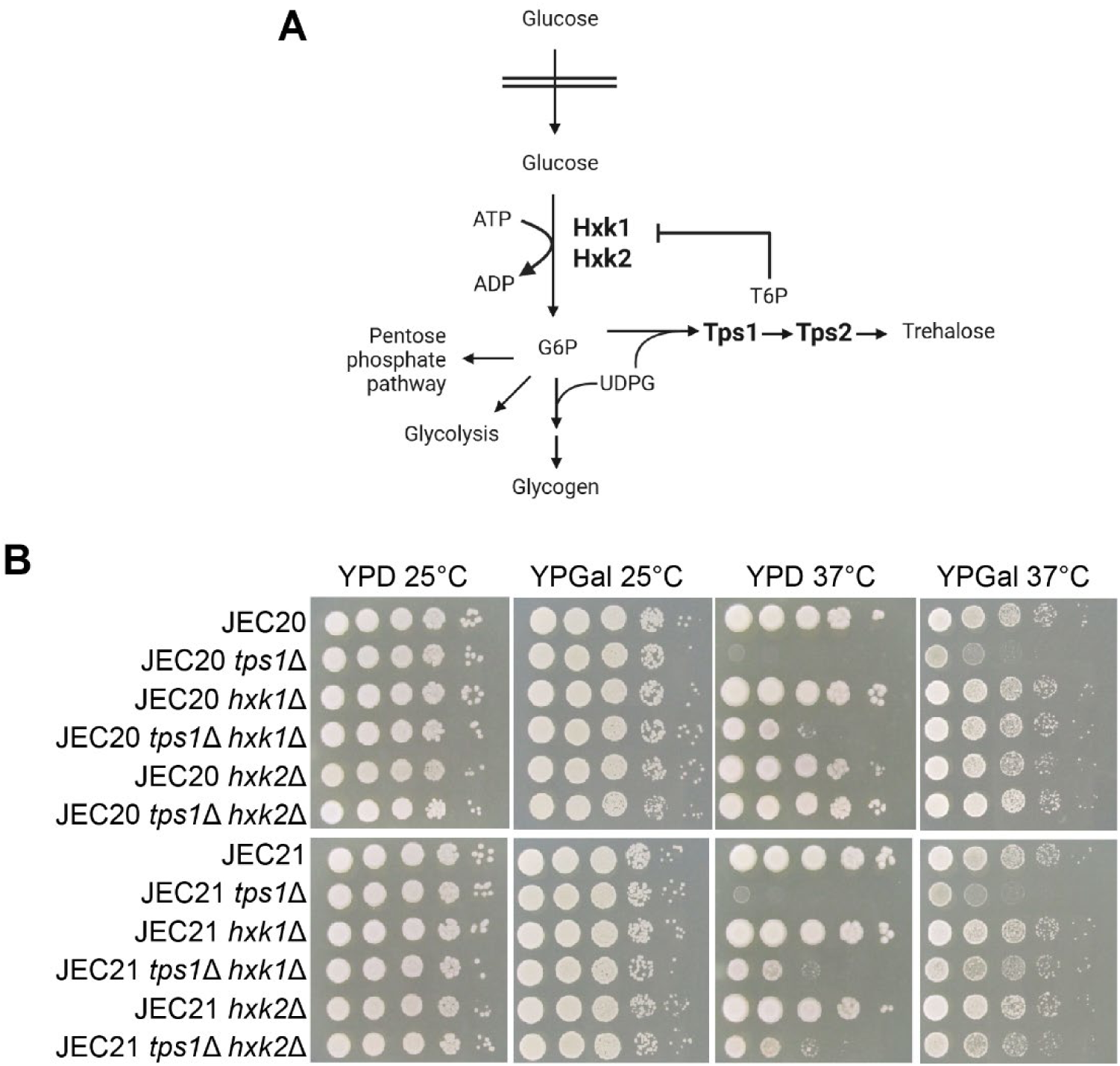
The connection between Tps1, hexokinases, and thermotolerance is glucose-dependent. **A)** A model for the integration of the trehalose biosynthesis pathway and phosphorylation of glucose mediated by hexokinases. After glucose enters the cells via glucose transporters, glucose is phosphorylated by the hexokinases Hxk1 and Hxk2. This is the first step in multiple energy-generating pathways. The product of Hxk1 and Hxk2 activity is glucose-6-phosphate (G6P). G6P is also a substrate of the trehalose biosynthesis enzyme, Tps1. Tps1 generates a signaling intermediate, trehalose-6-phosphate (T6P), which in several yeasts inhibits the activity of hexokinases. G6P is also an initiator of energy-generating processes, one of which leads to the production of glycogen. Drawn with Biorender. **B)** Serial dilution assays testing the ability of *C. deneoformans* wildtype and *tps1*Δ, *hxk1*Δ, *hxk2*Δ, *tps1*Δ *hxk1*Δ, and *tps1*Δ *hxk2*Δ strains to grow on YPD and YPGal at either 25°C or 37°C.

To further study the role of hexokinases in Tps1-dependent thermotolerance, we generated the following deletion strains in the *C. deneoformans* JEC20 and JEC21 congenic backgrounds: *tps1*Δ, *hxk1*Δ, *hxk2*Δ, *tps1*Δ *hxk1*Δ, and *tps1*Δ *hxk2*Δ. We performed serial dilution assays to investigate the growth of these strains at 25°C and 37°C on YPD and YPGal media (Figure 4B). JEC20 *tps1*Δ and JEC21 *tps1*Δ displayed a temperature-sensitive phenotype on YPD plates exposed to 37°C, as expected. The temperature-sensitive phenotype of JEC20 *tps1*Δ and JEC21 *tps1*Δ was partially rescued by growth on YPGal. JEC20 *hxk1*Δ and JEC21 *hxk1*Δ had no defect in thermotolerance, similar to the corresponding mutants in *C. neoformans* (47). However, deletion of *HXK1* in the *tps1*Δ mutants led to partial restoration of growth at 37°C on YPD medium. In contrast, JEC20 *hxk1*Δ *tps1*Δ and JEC21 *hxk1*Δ *tps1*Δ exhibited wild-type levels of growth on YPGal at 37°C. The greater restoration of growth on YPGal compared to YPD suggests that glucose acts as critical initiating factor in this pathway.

Although we did not detect a suppressor strain with a single mutation in the *HXK2* gene, as we did for *HXK1*, we tested the ability of deletion of *HXK2* to bypass *tps1* for thermotolerance (Figure 4B). JEC20 *hxk2*Δ and JEC21 *hxk2*Δ grow like wildtype at 37°C. Thus, *HXK2* is also critical for Tps1-mediated thermotolerance, as was demonstrated by the restored growth of *tps1*Δ *hxk2*Δ strains at 37°C grown on YPD plates, compared to *tps1*Δ strains. We also tested the growth of *tps1*Δ, *hxk1*Δ, *hxk2*Δ, *tps1*Δ *hxk1*Δ, and *tps1*Δ *hxk2*Δ in yeast-peptone (YP) media in the absence of a sugar source. Overall growth of the cells decreased on this media, however, the partial rescue of *tps1*Δ by deletion of *HXK1* or *HXK2* was still observed (Supplemental Figure 5).

To assess the ability of *C. deneoformans hxk1* and *hxk2* mutants to suppress the *tps1* mutant thermosensitivity in liquid cultures, growth assays were conducted at 25° and 37°C in both YPD and YPGal media (Supplemental Figure 6). All cultures grew well at 25°C. JEC20 *tps1*Δ displayed a temperature-sensitive phenotype in YPD cultures grown at 37°C. The *hxk2*Δ mutant displayed a slight growth defect in YPD when grown at 37°C, which may explain the paucity of spontaneous mutant isolates in *HXK2* in the suppressor screen. The *tps1*Δ growth defect at 37°C could be rescued by deletion of *HXK1* or *HXK2* in YPD, similar to the growth on solid medium. There was also a slight defect in the growth of JEC20 *tps1*Δ in YPGal at 37°C in the liquid growth curves. The *tps1*Δ growth defect in YPGal at 37°C was rescued by deletion of *HXK1* or *HXK2*.

### Suppressor mutations demonstrate alterations in glycolytic metabolites

To determine how the deletion of *C. deneoformans HXK1* or *HXK2* in the *tps1*Δ background affects glycolytic flux, we measured G6P accumulation in wild-type and mutant strains (Figure 5A). Deletion of *TPS1* resulted in a statistically significant 2-fold increase in G6P compared to the wildtype strain. Deletion of either *HXK1* or *HXK2* reduced the amounts of G6P to less than wild-type levels but with no clear differences in the levels of G6P between the *hxk1*Δ and *hxk2*Δ strains, indicating that they can still both phosphorylate glucose. In the *tps1*Δ *hxk1*Δ and *tps1*Δ *hxk2*Δ strains, G6P levels were reduced by approximately 20 – 40% relative to the corresponding *tps1*Δ strains, indicating that deletion of *HXK1* or *HXK2* in the *tps1Δ* background reduced the high levels of G6P that are likely due to derepression of the hexokinase activity. Interestingly, temperature did not affect the relative differences in G6P accumulation amongst the mutants.

**Figure 5.**
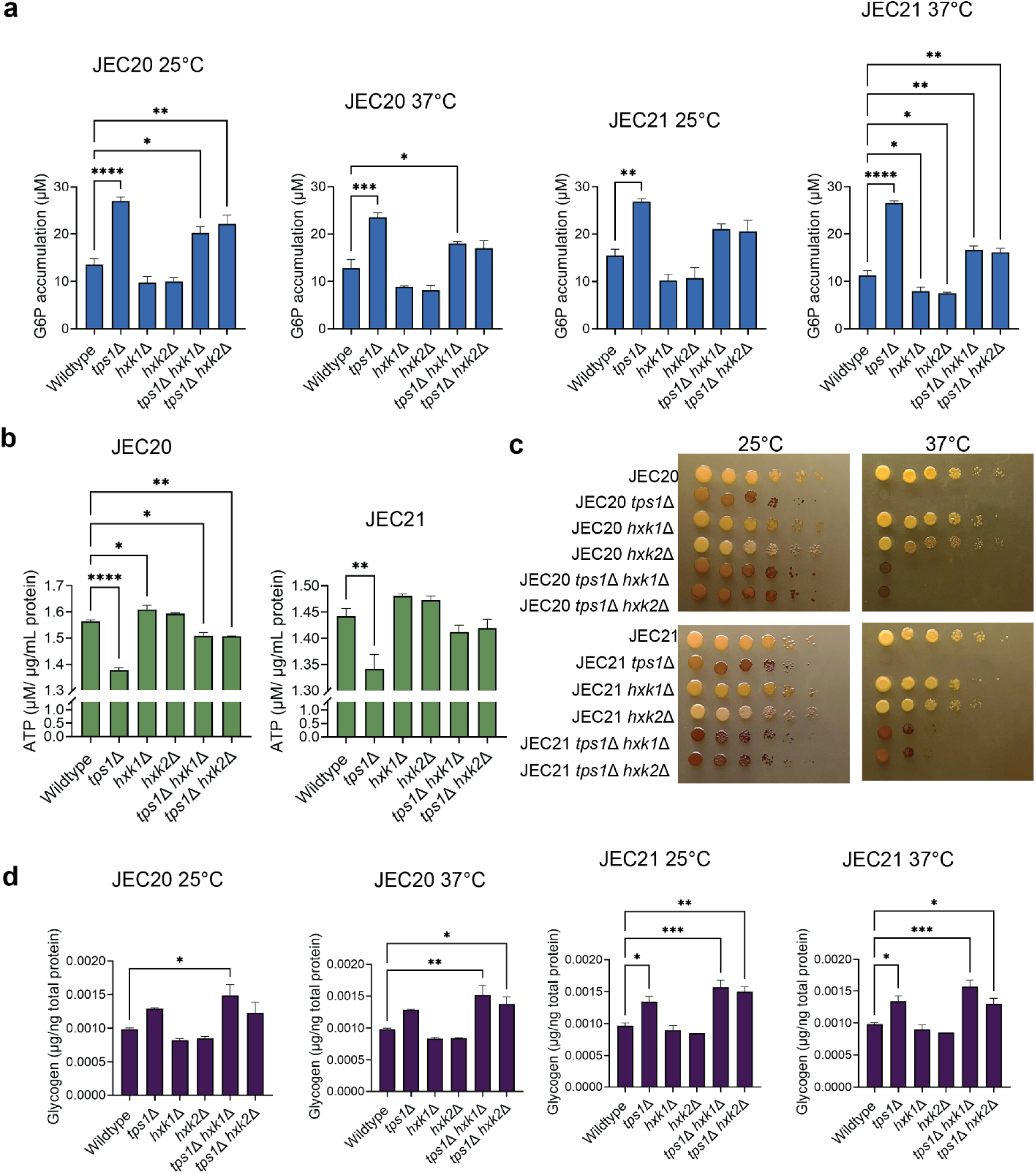
G6P, glycogen, and trehalose accumulation levels controlled by Tps1, Hxk1, and Hxk2. **A)** Measurement of G6P accumulation via an enzymatic-coupled and colorimetric assay in *C. deneoformans* wildtype and *tps1*Δ, *hxk1*Δ, *hxk2*Δ, *tps1*Δ *hxk1*Δ, and *tps1*Δ *hxk2*Δ strains in the JEC20 and JEC21 backgrounds grown in YPD liquid cultures at either 25°C or 37°C. Bars signify the mean ± SEM of biological triplicates. Data shown are representative of three independent experiments. **B)** Measurement of ATP concentration via an enzymatic-coupled and colorimetric assay in *C. deneoformans* wildtype and *tps1*Δ, *hxk1*Δ, *hxk2*Δ, *tps1*Δ *hxk1*Δ, and *tps1*Δ *hxk2*Δ strains in the JEC20 and JEC21 backgrounds grown in YPD liquid cultures at 25°C and exposed to 37°C for 1 minute. Bars signify the mean ± SEM of biological triplicates. Data shown are representative of three independent experiments. **C)** Iodine-stained serial dilution culture assays of *C. deneoformans* wildtype and *tps1*Δ, *hxk1*Δ, *hxk2*Δ, *tps1*Δ *hxk2*Δ, and *tps1*Δ *hxk2*Δ strains to assess glycogen accumulation at 25°C or 37°C. **D)** Measurement of glycogen accumulation via an enzymatic-coupled and colorimetric assay in *C. deneoformans* wildtype and *tps1*Δ, *hxk1*Δ, *hxk2*Δ, *tps1*Δ *hxk2*Δ, and *tps1*Δ *hxk2*Δ strains grown at 25°C or 37°C. Bars signify the mean ± SEM of biological triplicates. Data shown are representative of three independent experiments.

To determine how deletion of *C. deneoformans HXK1* and *HXK2* in the *tps1*Δ background affects ATP expenditure, we then measured ATP levels in wild-type and mutant strains after the cells were exposed to heat shock at 37°C for 1 minute (Figure 5B). Deletion of *TPS1* resulted in a significant decrease (approximately 10%) in ATP compared to the wild-type strain at 37°C. This is consistent with the prior observation with the rapid decrease in ATP levels found in *C. albicans tps1*Δ/*tps1*Δ at 42°C (48). Furthermore, deletion of either *HXK1* or *HXK2* increased the concentration of ATP to slightly more than wild-type levels. However, there were no clear differences in the levels of ATP between the *hxk1*Δ and *hxk2*Δ strains, indicating that they can both utilize ATP as a substrate, as expected. In the *tps1*Δ *hxk1*Δ and *tps1*Δ *hxk2*Δ strains an intermediate level of ATP was detected. Therefore, deletion of *HXK1* or *HXK2* in the *tps1*Δ background increased the depleted levels of ATP at 37°C that are likely due to the derepression of the hexokinase activity.

The storage carbohydrate glycogen, of which G6P and UDP-glucose are precursors, also accumulated in the *tps1*Δ strains (Figure 5C,D). Interestingly, the *tps1*Δ *hxk1*Δ and *tps1*Δ *hxk2*Δ strains displayed levels of glycogen similar to the *tps1*Δ levels (Figure 5C,D).

Combined, these data provide evidence that the increase in hexokinase activity and subsequent overaccumulation of G6P and depletion of ATP levels contribute to the temperature sensitivity of *C. deneoformans tps1*Δ mutants. Deletion of *HXK1* or *HXK2* in the *tps1*Δ mutant background partially restores the growth at elevated temperatures by rebalancing glycolytic flux to near wild-type levels, as reflected by G6P and ATP levels. In contrast, glycogen remains hyperaccumulated and is not corrected by deletion of *HXK1* or *HXK2*.

## Discussion

In this study, we conducted a genetic suppressor screen to isolate spontaneous thermotolerant mutants in the *C. deneoformans tps1*Δ background, which is inviable at 37°C (Figure 1A). The predominant finding from the whole-genome sequencing is the occurrence of loss-of-function mutations in the *HXK1* gene in 17 of the 21 *tps1*Δ suppressors (Table 1). Further, we also isolated a suppressor strain with a mutation in the gene *HXK2* encoding the additional hexokinase 2 in *C. deneoformans* in combination with a mutation in *SNF5*, a component of the SWI/SNF chromatin-remodeling complex (Table 1). We also identified other suppressor mutations in genes associated with various cellular pathways, including an arrestin protein or a retrograde cargo transporter. One such factor, Ali1, an arrestin-domain protein, was identified in the suppressor screen and may influence thermotolerance through effects of cellular organization and trafficking. However, its specific contribution remains to be determined (49, 50). Here, we investigated the finding that the ability of a *C. deneoformans tps1*Δ mutant to actually tolerate elevated temperatures is increased by loss-of-function mutations in *HXK1*.

The Hxk1 point mutations identified in the suppressors span locations across the protein from the N-terminal domain to both the small and large domains (Figure 2B,C). While none of these mutations have been implicated directly in substrate-binding or catalysis, they are largely conserved by identity or with similarity and thus, are likely to have key structural roles in Hxk1 activity (Supplemental Figure 2). We chose to investigate the function of hexokinase variants that will provide information regarding the contribution of understudied regions of the protein to hexokinase function. Structural predictions did not reveal how these point mutations affect the overall hexokinase structure (Supplemental Figure 4). However, we could detect a decrease in the confidence of the structural model by mutating G537 to cysteine, based on the pLDDT score, indicating the importance of this residue that is proximal to the ATP-binding site (Supplemental Figure 4). Hxk1 W253S is mutated on the small domain near the ATP-binding site and may affect the binding of ATP to hexokinase (Figure 2C). In fact, most interesting, is the mutation of Hxk1 G388 to aspartic acid, which is not proximal to any known substrate-binding sites (Figure 2C). Introduction of aspartic acid may result in global changes that affect the function of hexokinases allosterically. It has been previously shown before that allosteric regulation is important to the function of Hxk2 in *S. cerevisiae* (51). Additionally, G388 is predicted to be near the relatively flexible N-terminal domain of Hxk1 and this may also indicate some functional relevance of the domain as well. Despite the multiple locations of the point mutations in this protein, we demonstrated that these alterations led to reductions in Hxk1 activity measured both *in vitro* with recombinant proteins expressed in *E. coli* and further from *C. deneoformans* cell extracts (Figure 3). Future work using both biochemical and structural approaches will help determine the specific role of these point mutations on the function of Hxk1.

The enzymatic function of hexokinases is to phosphorylate glucose following the entry of glucose into the cell. This reaction yields G6P, which then goes proceeds to initiate multiple energy-generating pathways, including both glycolytic and pentose-phosphate pathways (32, 33). Thus, hexokinase activity is critical for cellular metabolism and yeast survival under conditions in which glucose is a major energy source. In *S. cerevisiae* and other fungal organisms, hexokinase activity is regulated by T6P, the product of the first enzyme in the trehalose biosynthesis pathway, Tps1 (43–46). T6P competitively inhibits the activity of hexokinase (44). The inhibition of hexokinase by T6P regulates the entry of glucose into glycolysis. As a result, glucose-containing media are toxic for *tps1*Δ mutants, whereas growth of *tps1*Δ mutants on other media, such as YPGal, is improved (19, 52–54). Van Heerden *et*. *al*., described the lack of inhibition of hexokinases as resulting in an imbalanced glycolytic state, in which ATP consumption in the beginning of glycolysis outpaces ATP production in the later stages of glycolysis (55). Failure to maintain balance between the two stages of glycolysis results in a dysfunctional cellular state and ultimately results in a loss of cell viability (55). In the unbalanced glycolytic state, levels of metabolites central to the initial stages of glycolysis, including ATP and inorganic phosphate, escape normal regulatory constraints. The contribution of the trehalose biosynthesis pathway in yeast, in combination with the upper portion of the glycolytic pathway, forms a biochemical cycle to tune the dynamics of glycolysis towards a functional metabolic state (55). Our results revealed alterations in the levels of metabolites surrounding the entry point into glycolysis, G6P and ATP, and a further downstream product, glycogen. G6P and ATP levels are disrupted in the *tps1*Δ mutant and are partially restored to wild-type levels by deletion of either *HXK1* or *HXK2,* whereas glycogen remains elevated in the double mutants (Figure 5). ATP levels are depleted in *tps1*Δ mutants after heat shock at 37°C (48). Maintenance of proper ATP levels in cells is critical for viability. Therefore, deletion of either *HXK1* or *HXK2* in a *tps1*Δ mutant enables the restoration of ATP levels after exposure to elevated temperatures. Glycogen, an energy storage polymer that is generated in a pathway downstream of glycolysis, hyperaccumulates in the *tps1*Δ mutant, providing more evidence of unbridled glucose metabolism in *tps1*Δ mutants (56–58). This hyperaccumulation persists in *tps1*Δ *hxk1*Δ and *tps1*Δ *hxk2*Δ mutants, likely reflecting an expanded UDP-glucose pool that is preferentially directed toward glycogen synthesis in the absence of Tps1. These data reveal a mechanism by which Tps1 regulates glycolytic flux to promote thermotolerance in *C. deneoformans* (Figure 6).

**Figure 6.**
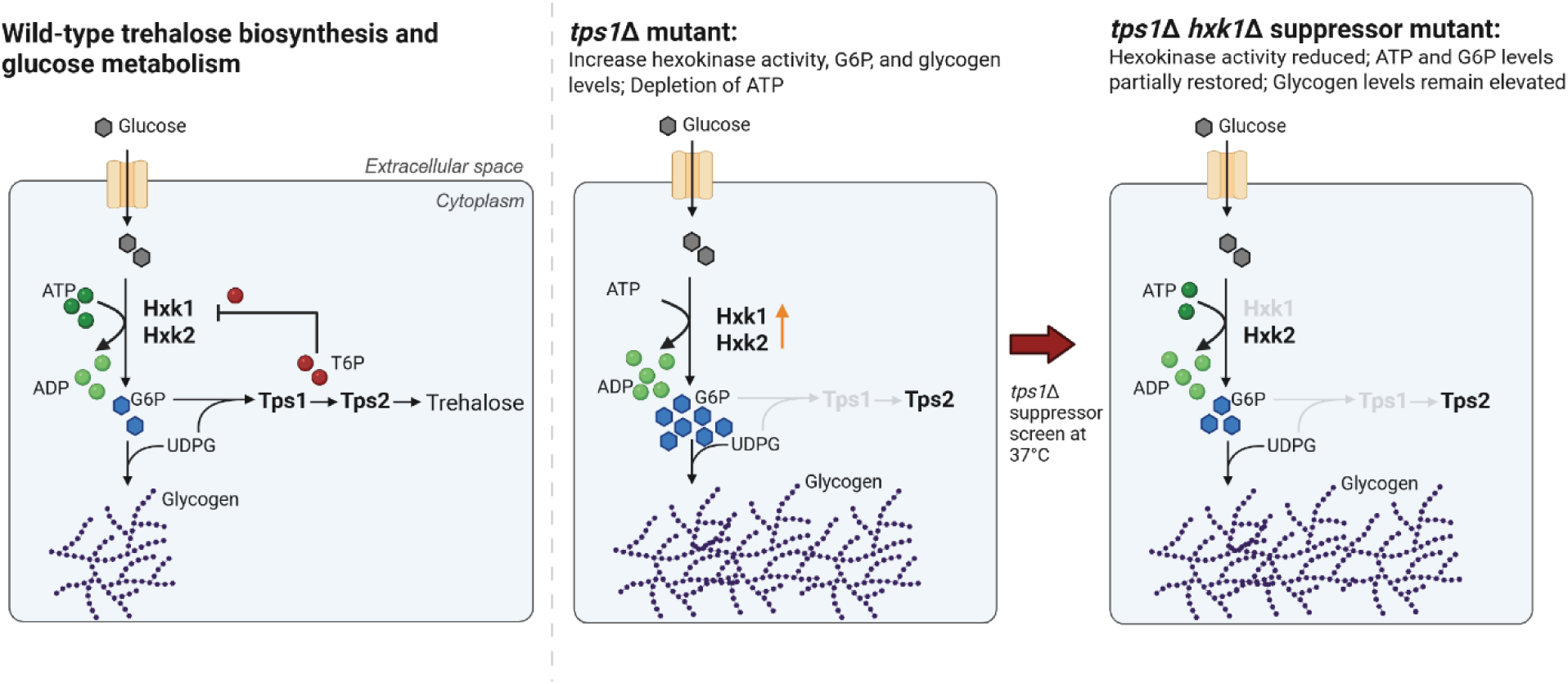
A model depicting how Tps1 governs glycolytic flux in *C. deneoformans* to promote thermotolerance. In wild-type *C. deneoformans* cells, after glucose enters the cells via glucose transporters, glucose is phosphorylated by the hexokinases Hxk1 and Hxk2. This step requires the expenditure of ATP and leads to the generation of glucose-6-phosphate (G6P). G6P is a substrate of the trehalose biosynthesis enzyme, Tps1, and also a precursor for glycogen, a glucose storage molecule. Tps1 generates a signaling intermediate, trehalose-6-phosphate (T6P), an inhibitor of hexokinases. In *C. deneoformans tps1*Δ mutants, which are unable to survive at elevated temperatures, the repression of T6P is removed. This loss of repression allows an increase in hexokinase activity, depletion of ATP, and a hyperaccumulation of G6P and glycogen. Disruption of hexokinase activity enables *C. deneoformans tps1*Δ mutants to survive at 37°C, due to a reduction in hexokinase activity and a partial restoration of ATP and G6P to wild-type levels, while glycogen levels are maintained at elevated levels. Drawn with Biorender.

Interestingly, we found that the unbalanced glycolytic state in the *tps1*Δ mutant exists at both 25°C and 37°C (Figure 5). Furthermore, alterations in G6P levels in the *tps1*Δ, *hxk1*Δ, *hxk2*Δ, *tps1*Δ *hxk1*Δ, and *tps1*Δ *hxk2*Δ mutant backgrounds occurred in a similar pattern at both 25°C and 37°C. Therefore, we posit that at higher temperatures, the heat-induced stress adds additional strain to already stressed cellular systems. A stressed *tps1*Δ mutant, with unregulated glycolytic flux may have less capacity to tolerate elevated temperatures, which then may cause membrane instability, protein misfolding, and increased formation of reactive oxygen species, as well as, result in a lack of protection of the cell surface and proteins by trehalose (59, 60).

Trehalose is essential for stress response, and in *Cryptococcus* species, it regulates growth at 37°C, which is crucial for virulence. Here, we showed this conserved role in *C. deneoformans*, in addition to *C. neoformans* and *C. gattii* that have been previously described. Additionally, the relationship between Tps1 and Hxk1 that we found was also previously identified in *C. gattii,* where deletion of either *HXK1* or *HXK2* in *tps1*Δ suppressed the temperature-sensitive phenotype of *C. gattii tps1*Δ (18), leading to the hypothesis that the regulation of glycolysis via the trehalose biosynthesis pathway is critical for yeast growth at 37°C. Our results in this study indeed showed that the trehalose biosynthesis pathway is crucial for regulating glycolytic flux, hinting at the conservation of this pathway between these two species. Whether this conservation extends to the third pathogenic species in this complex, *C. neoformans*, remains to be investigated. Interestingly, our efforts to recover spontaneous suppressor isolates for growth at 37°C employing a *C. neoformans tps1*Δ mutant failed, suggesting more tight and differential regulation of this pathway. Beyond the *Cryptococcus* pathogenic species complex, a similar relationship between trehalose biosynthesis and glycolysis has been described in other model and pathogenic fungi of plants and humans. In *S. cerevisiae* and plant pathogen *Magnaporthe grisea* genetic disruption of hexokinase has been shown to partially restore metabolic and growth defects associated with loss of Tps1/Tps2 function (43, 44, 61). Early work in *S. cerevisiae* demonstrated that deletion of *TPS1*(*GGS1*) results in uncontrolled influx into glycolysis, accumulation of sugar phosphates, and impaired growth on fermentable carbon sources (46). Petzold *et*. *al*., postulated the poor growth of *tps1*Δ mutants at 37°C on glucose could be due to the loss of regulation of glycolysis, leading to the increase in sugar phosphates and decrease in ATP production, consistent with our findings presented here (19). Interestingly, deletion of *HXK1* in the *M. grisea tps1*Δ mutant resulted in partial restoration of growth in the presence of glucose but did not suppress virulence or sporulation phenotypes. A recent study in the human pathogen *Candidozyma auris* showed that deletion of *TPS1* results in a cell wall defect and altered antifungal drug resistance due to inhibition of hexokinase activity via accumulated T6P, linking the two pathways (62). Further studies will be required to assess such impacts as well as restoration of virulence by *hxk1* deletion in *C. deneoformans tps1*Δ mutants.

Biochemical and structural studies will be necessary to address certain unanswered questions in this system, such as how does T6P competitively inhibit the activity of *C. deneoformans* Hxk1 and Hxk2? We speculate that the binding of T6P may affect the binding of native substrates, ATP and glucose, as well as the release of G6P, which is necessary for the catalytic process. Conformational changes are critical to the catalytic function of hexokinases. The binding of T6P may “lock” hexokinases in an inactive conformation. Identification of novel scaffolds and mechanisms to disrupt hexokinase activity, such as T6P, is important, given that hexokinases are major targets of cancer drugs (63, 64). Although humans lack the trehalose biosynthesis pathway, previous studies have demonstrated that T6P can specifically inhibit human HK2 (hexokinase 2) but not human HK1 (hexokinase 1) (65).

In total, this work provides a detailed mechanistic study of the *C. deneoformans* dependency on Tps1 for cryptococcal thermotolerance. We identified genetic mutations that bypass the requirement of Tps1 for *C. deneoformans* survival at 37°C. Furthermore, we conclude that regulation of glycolytic flux plays a key role in the requirement for Tps1, a trehalose biosynthesis enzyme, for high temperature growth in *C. deneoformans*, highlighting significant ways in which temperature-sensitive fungi may evolve when exposed to increasing environmental temperatures. As global temperatures rise, evolutionary adaptations that allow fungi such as *C. deneoformans* to thrive at higher temperatures may become increasingly relevant. The same adaptations that permit survival in a warming environment may concurrently promote the emergence of new fungal pathogens.

## Data Availability

The authors affirm that all representative data necessary for confirming the conclusions of the article are present within the article, figures, and tables. Strains and primers used in this work are listed in Supplementary Tables 1 and 2, respectively, and are available upon request. The whole genome sequencing data is available at NCBI under the BioProject accession number PRJNA1348356. The DNA sequences of codon-optimized *C. deneoformans* Hxk1 expression constructs, including the wild-type sequence and four variants used in this study, have been deposited in NCBI GenBank as synthetic constructs under accession numbers PZ354051, PZ354052, PZ354053, PZ354054, and PZ354055.

## Acknowledgments

We thank Dr. Marc Meneghini for insights into the role of trehalose in cellular adaptation to changing glucose levels in *S. cerevisiae*. We are grateful to Dr. David Hubert, Dr. Andrew Alspaugh, Dr. Zoë Hilbert, and Dr. John Perfect for their helpful discussions, suggestions, and comments on the manuscript. We thank Dr. Alejandro Antonia of the Alspaugh laboratory for advice on glycogen assays. We thank Dr. Devjanee (Devi) Swain Lenz and the Duke University School of Medicine for the use of the Sequencing and Genomic Technologies (SGT) Shared Resource, which provided whole-genome sequencing via the Illumina Novaseq platform. We thank Dr. Xiaorong Lin for generously providing additional strains used in this study.

## Author Contributions

V.Y. and E.J.W. conducted most of the experiments and interpreted data and results. K.A.C. conducted some experiments. V.Y. and E.J.W. wrote the original manuscript draft of the manuscript. V.Y., J.H., and E.J.W. conceived the study and interpreted data and results. V.Y., J.H., and E.J.W. edited the manuscript.

