## Supplemental Material for "Trehalose-6-phosphate synthase promotes thermotolerance by governing glycolytic flux in *Cryptococcus deneoformans*"

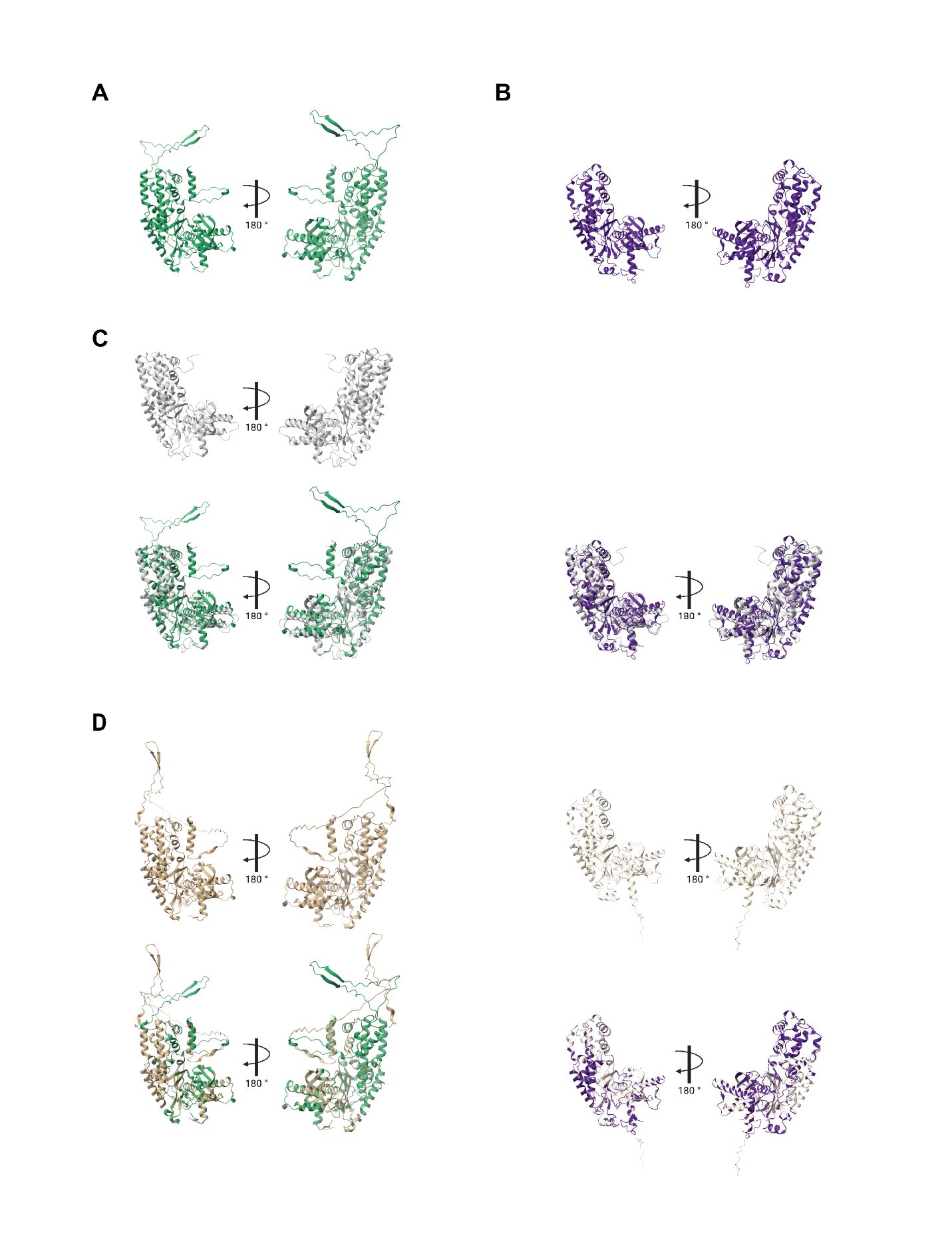

**Supplemental Figure 1. Predicted structures of *C. deneoformans* Hxk1 and Hxk2 reveal high structural conservation with other yeast hexokinase homologs. A)** AlphaFold-predicted structure of *C. deneoformans* Hxk1 (AlphaFold Q5KCL1) shown in green. **B)** Alpha-Fold predicted structure of *C. deneoformans* Hxk2 (AlphaFold Q5KM78) shown in purple. **C)** Structural overlays of *C. deneoformans* hexokinases with *K. lactis* Hxk1 (PDB: 3O1W). The top panels show *K. lactis* Hxk1 alone (grey), and the bottom panels show structural alignments with *C. deneoformans* Hxk1 (green, left) and *C. deneoformans* Hxk2 (purple, right). **D)** Structural overlays of *C. deneoformans* Hxk1 and Hxk2 with *C. neoformans* var. *grubii* H99 hexokinase homologs. The top panels show AlphaFold-predicted structures of *C. neoformans* Hxk1 (J9VWU8, brown, left) and Hxk2 (J9VNK3, beige, right) alone. The bottom panels show structural alignments of *C. deneoformans* Hxk1 (green) with *C. neoformans* Hxk1 (brown, left) and *C. deneoformans* Hxk2 (purple) with *C. neoformans* Hxk2 (beige, right). Paired views are related by a 180% rotation around the y-axis.

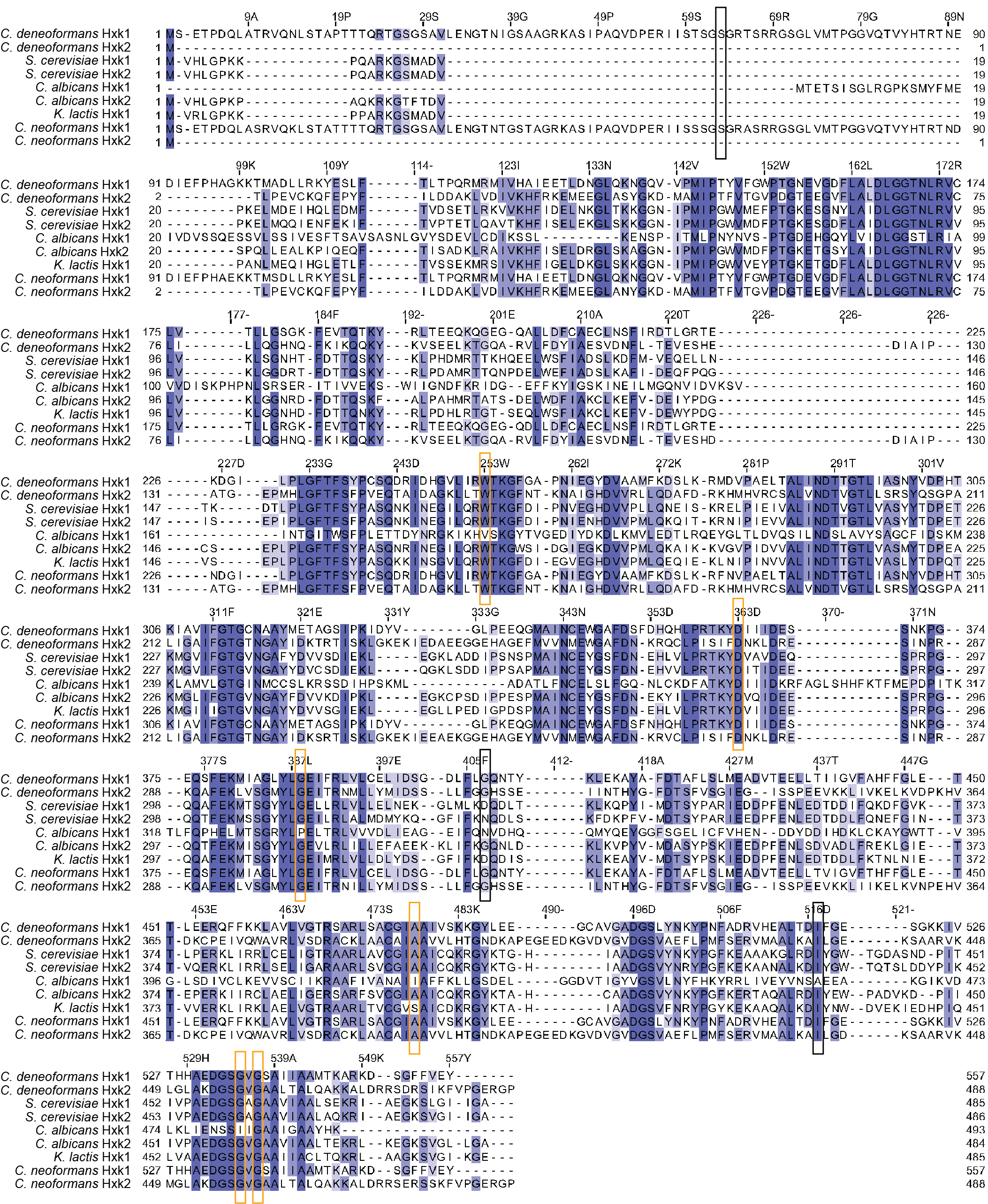

**Supplemental Figure 2. Hexokinase protein sequence alignment reveals conservation.** Clustal Omega was used to align the following hexokinases sequences: *C. deneoformans* JEC21α Hxk1 (CNH01400), *C. deneoformans* JEC21α Hxk2 (CNB02660), *S. cerevisiae* Hxk1 (KZV11659), *S. cerevisiae Hxk2* (CAA96973), *C. albicans Hxk1* (AOW30374), *C. albicans Hxk2* (AOW31187), *K. lactis* Hxk1 (QEU60788), *C. neoformans* var. *grubii* H99 Hxk1 (CNAG_05480), and *C. neoformans* var. *grubii* H99 Hxk2 (CNAG_03769) (1). Orange rectangles highlight the *C. deneoformans* Hxk1 residues that were identified in the suppressor screen. Black rectangles highlight the *C. deneoformans* Hxk1 residues that are modified due to either a nonsense mutation or frameshift mutation. The residues in blue are 50% or more conserved among the hexokinase sequences. The alignment is visualized using Jalview (2).

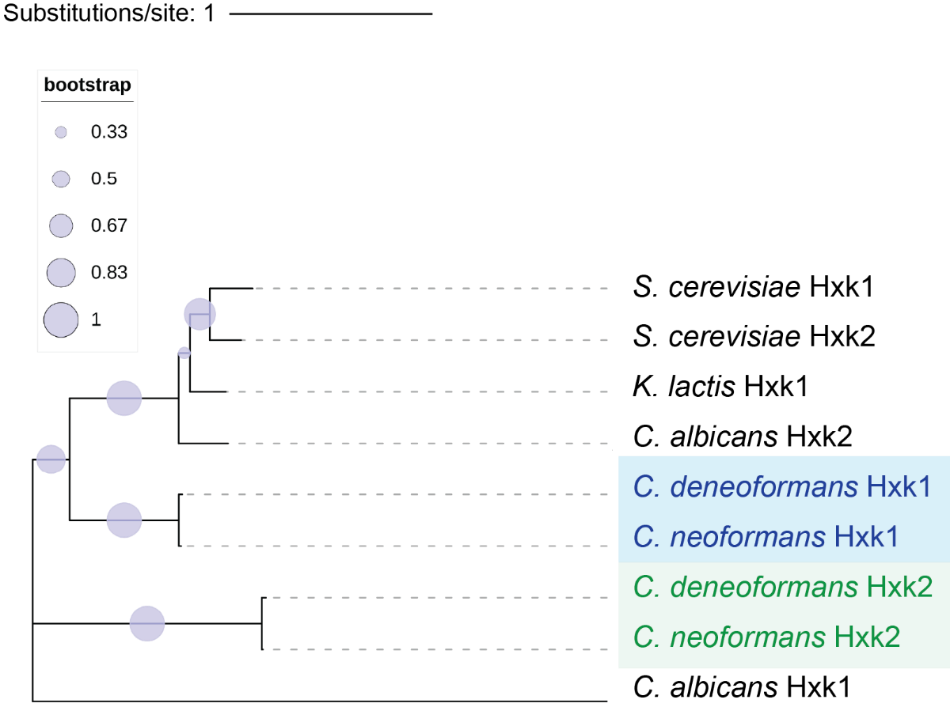

**Supplemental Figure 3. Phylogenetic analysis of hexokinase orthologs across various fungal species reveals clustering of *Cryptococcus* hexokinases.** The phylogeny was inferred using the Maximum Likelihood method and the Le-Gascuel LG model (3). The bootstrap consensus tree inferred from 100 replicates is taken to represent the evolutionary history of the taxa analyzed where branches corresponding to partitions reproduced in less than 50% of replicate trees are collapsed (4). The percentage of replicate trees in which the associated taxa clustered together (100 replicates) is shown next to the branches (4). Initial trees for the heuristic search were obtained by applying the Neighbor-Joining method to a matrix of pairwise distances estimated using p-distance (5). The evolutionary rate differences among sites were modeled using a discrete Gamma distribution across 5 categories, with 8.11% of sites deemed evolutionarily invariant. The analytical procedure encompassed 9 amino acid sequences with 629 positions in the final dataset. Evolutionary analyses were conducted in MEGA12 (6). Tree is illustrated using iTol software (7).

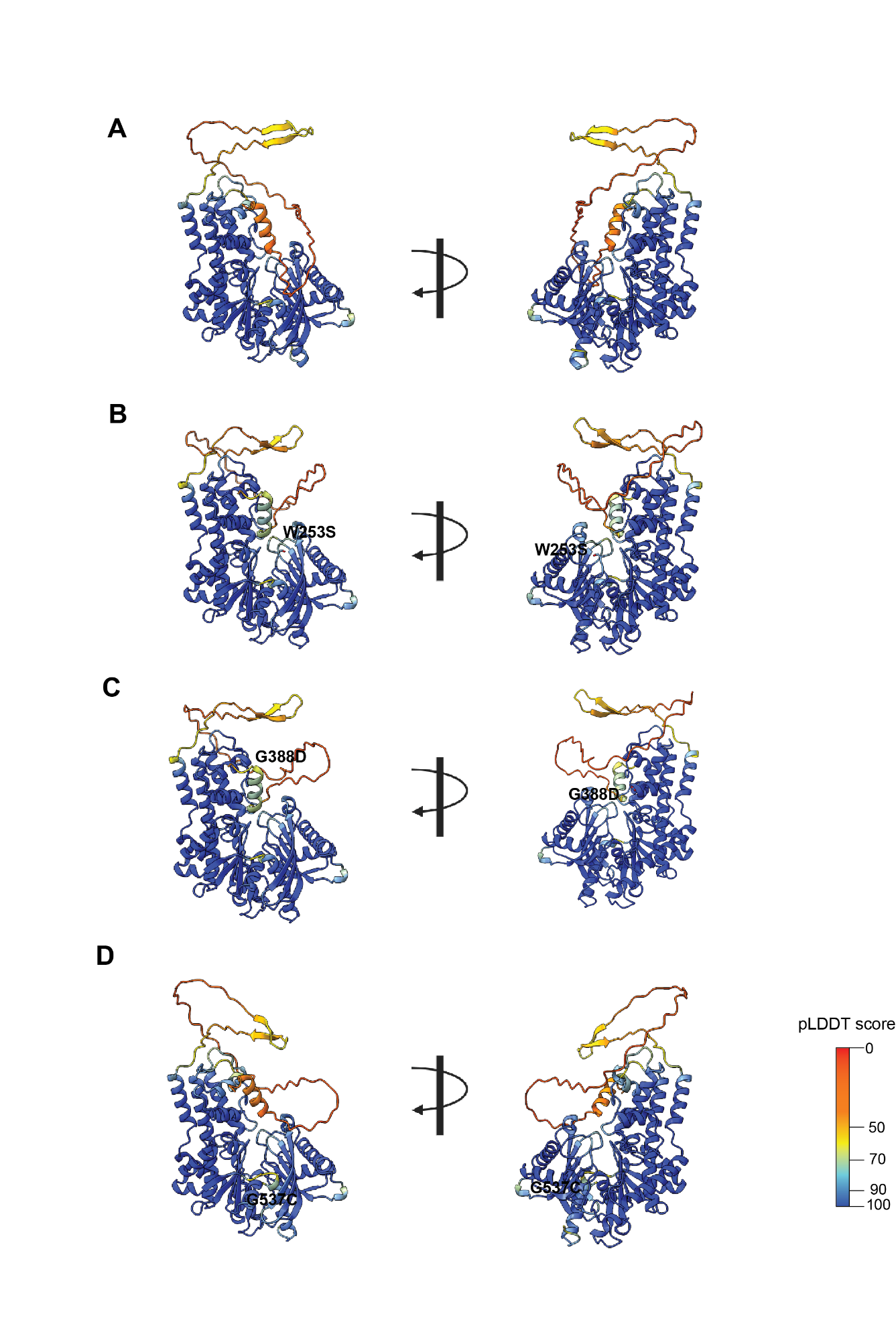
**Supplemental Figure 4. AlphaFold predictions of Hxk1 variant structures.** Shown in ribbon diagrams are the AlphaFold predictions (8) of wild-type *C. deneoformans* Hxk1 (**a**) and the Hxk1 variants W253S (**b**), G388D (**c**), and G537C (**d**). The mutant residues are shown in heteroatom-colored stick representation. The structures are displayed using the continuous pLDDT color scale applied by the AlphaFold Colab pipeline, ranging from red/orange (low confidence) through yellow to cyan/blue (high confidence). Figures were visualized and rendered using UCSF ChimeraX (9).

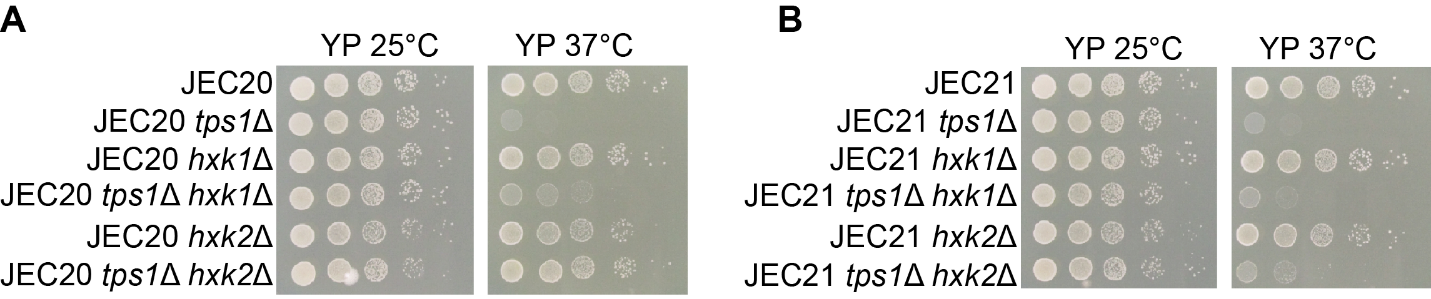

**Supplemental Figure 5. Growth of strains on yeast extract-peptone media lacking glucose.** **a**) *C. deneoformans* JEC20**a** wildtype, *tps1*Δ, *hxk1*Δ, *tps1*Δ *hxk1*Δ, *hxk2*Δ, and *tps1*Δ *hxk2*Δ mutant strains were serially diluted and grown on yeast-peptone media at 25°C and 37°C. **b**) *C. deneoformans* JEC21α wildtype, *tps1*Δ, *hxk1*Δ, *tps1*Δ *hxk1*Δ, *hxk2*Δ, and *tps1*Δ *hxk2*Δ mutant strains were serially diluted and grown on yeast-peptone media at 25°C and 37°C.

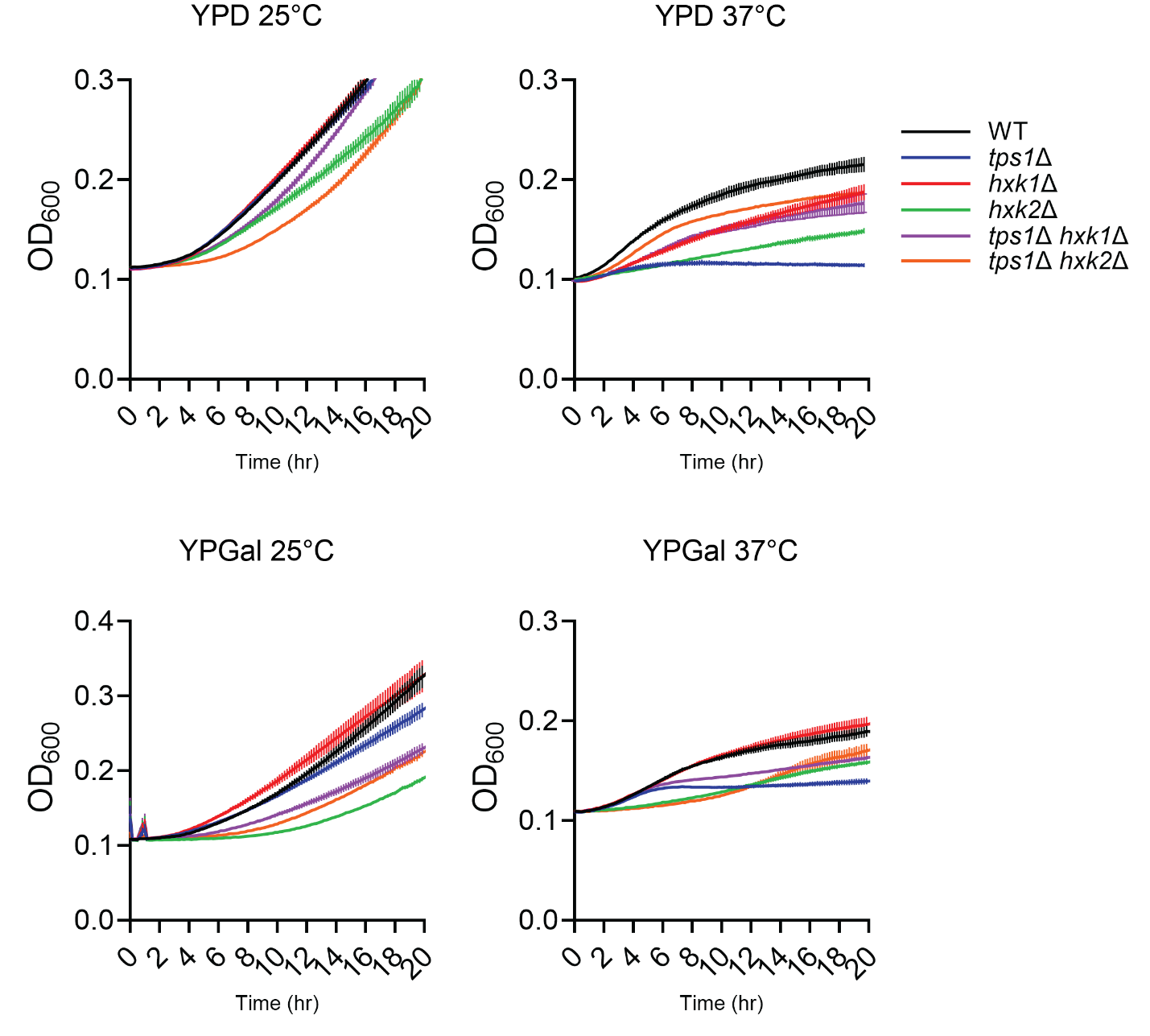

**Supplemental Figure 6. Deletion of *HXK1* or *HXK2* rescues the growth defect of tps1 in liquid media.** Growth curve analysis of *C. deneoformans* JEC20**a** wildtype and *tps1*Δ, *hxk1*Δ, *tps1*Δ *hxk1*Δ, *hxk2*Δ, and *tps1*Δ *hxk2*Δ mutant strains in YPD or YPGal media grown at either 25°C and 37°C. Optical density at 600 nm was measured every 10 minutes for 20 hours. Color legends for the different strains are given. Error bars represent the standard error of the mean (SEM). Graphs are representative of three biological replicates.

**Supplemental Tables**

**Table S1. List of strains used in this study.**

| Strain name | Description | Reference |
| --- | --- | --- |
| JEC21α | Wild-type *MAT*α | (10) |
| JEC20a | Wild-type *MAT***a** | (10) |
| XL468 | JEC20**a** *tps1*Δ | (11) |
| XL470 | JEC21α *tps1*Δ | (11) |
| VYD489 | JEC20a *tps1*Δ*::NAT CNH01400*_G388D | This study |
| VYD490 | JEC20a *tps1*Δ*::NAT CNH01400*_G388D | This study |
| VYD492 | JEC20a *tps1*Δ*::NAT CNH01400*_G388D | This study |
| VYD493 | JEC20a *tps1*Δ*::NAT CNH01400*_G407* | This study |
| VYD494 | JEC20a *tps1*Δ*::NAT CNH01400*_A478P | This study |
| VYD495 | JEC20a *tps1*Δ*::NAT CNB02660*_Frame shift at V420  JEC20a *tps1*Δ*::NAT CNA07190*_Q748* | This study |
| VYD496 | JEC20a *tps1*Δ*::NAT CNH01400*_gene deletion | This study |
| VYD497 | JEC20a *tps1*Δ*::NAT CNH01400*_W253S | This study |
| VYD498 | JEC20a *tps1*Δ*::NAT CNC04740*_Frame shift at N229 | This study |
| VYD499 | JEC20a *tps1*Δ*::NAT CNH01400*_gene deletion | This study |
| VYD500 | JEC20a *tps1*Δ*::NAT CNH01400*_Intron9 branch point | This study |
| VYD510 | JEC21α *tps1*Δ*::NAT CNH01400*_gene deletion | This study |
| VYD508 | JEC21α *tps1*Δ*::NAT CNC02740*_P1056Q | This study |
| VYD511 | JEC21α *tps1*Δ*::NAT CNH01400*_Intron2 splice donor  JEC21α *tps1*Δ*::NAT CNL04350*_Q12H | This study |
| VYD512 | JEC21α *tps1*Δ*::NAT CNC04740*_V299F | This study |
| VYD513 | JEC21α *tps1*Δ*::NAT CNH01400*_G535D | This study |
| VYD514 | JEC21α *tps1*Δ*::NAT CNH01400*_Frame shift at I517 | This study |
| VYD515 | JEC21α *tps1*Δ*::NAT CNH01400*_Intron 12 | This study |
| VYD516 | JEC21α *tps1*Δ*::NAT CNH01400*_G537C | This study |
| VYD517 | JEC21α *tps1*Δ*::NAT CNH01400*_Frame shift at S63 | This study |
| VYD518 | JEC21α *tps1*Δ*::NAT* *CNH01400*_D363N | This study |
| VYD607 | JEC20**a** *hxk1*Δ*::NEO* | This study |
| VYD608 | JEC20**a** *tps1*Δ*::NAT hxk1*Δ*::NEO* | This study |
| VYD609 | JEC20**a** *hxk2*Δ*::NEO* | This study |
| VYD610 | JEC20**a** *tps1*Δ*::NAT hxk2*Δ*::NEO* | This study |
| VYD611 | JEC21α *hxk1*Δ*::NEO* | This study |
| VYD612 | JEC21α *tps1*Δ*::NAT hxk1*Δ*::NEO* | This study |
| VYD613 | JEC21α *hxk2*Δ*::NEO* | This study |
| VYD614 | JEC21α *tps1*Δ*::NAT hxk2*Δ*::NEO* | This study |

***** denotes a nonsense mutation.

**Table S2. List of primers employed in this study.**

| Name | Sequence | Purpose |
| --- | --- | --- |
| JOHE55530 | TTGGAGGAGTGCTGAGTGGC | *HXK1* deletion construct |
| JOHE55531 | ATACCTCTCTTGCTGTTGGC |  |
| JOHE55532 | TCGTCGAATTTCGGCAACGGTC |  |
| JOHE55533 | CAGCTCACATCCTCGCAGCCATGACATGGCTGTGCGCTGG |  |
| JOHE55534 | CCAGCGCACAGCCATGTCATGGCTGCGAGGATGTGAGCTG |  |
| JOHE55535 | CCTTCCTGGCCTTGGTCATTCGGTTTATCTGTATTAACACGG |  |
| JOHE55536 | CCGTGTTAATACAGATAAACCGAATGACCAAGGCCAGGAAGG |  |
| JOHE55537 | TCCTGGAGTGTCATTTGCACC |  |
| JOHE55538 | ACCGGCAGGGTATACTGTTGCGGCGCCTACAACCACCCAGGTTTTAGAGCTAGAAATAGC | *HXK1* deletion guide RNA |
| JOHE55539 | ACCGGCAGGGTATACTGTTGCTGAACCATCCTCAGCATGGGTTTTAGAGCTAGAAATAGC |  |
| JOHE55874 | GATGAGGTTATTGAGAGCAACTCG | *HXK2* deletion construct |
| JOHE55875 | GTAAGCCTATCACGATATCAGTCC |  |
| JOHE55876 | TCATGTATATCCAAGGTTCTTG |  |
| JOHE55877 | CAGCTCACATCCTCGCAGCCATAGTGGTGTTATTTCGAGGCTC |  |
| JOHE55878 | GAGCCTCGAAATAACACCACTATGGCTGCGAGGATGTGAGCTG |  |
| JOHE55879 | GCCAGGGACGAACTTGATTGACGGTTTATCTGTATTAACACGG |  |
| JOHE55880 | CCGTGTTAATACAGATAAACCGTCAATCAAGTTCGTCCCTGGC |  |
| JOHE55881 | TAAGAAAGCAGTCGATAGCACC |  |
| JOHE55882 | ACCGGCAGGGTATACTGTTGGCACTTCCGTAAGGAGATGGGTTTTAGAGCTAGAAATAGC | *HXK2* deletion guide RNA |
| JOHE55883 | ACCGGCAGGGTATACTGTTGTGAGTGACTAGAAATCCTCGGTTTTAGAGCTAGAAATAGC |  |
| JOHE55941 | TCTGTCTCGTTACTCTTCTAGG | *HXK1* ORF PCR |
| JOHE55942 | GATAATGTCGTACTTGGTTCGG |  |
| JOHE55943 | CAACACGAAGAACGCTATCGG | *HXK2* ORF PCR |
| JOHE55944 | AATGGAGATAGGTAGGCACTGC |  |
